# ESCRT proteins organise stress-induced organelle contact between endoplasmic reticulum and Golgi

**DOI:** 10.64898/2026.05.29.728859

**Authors:** Oliver Pajonk, Lis Albert, Jasmin A. Schäfer, Leanne de Jager, Carlos Martín de Hijas, Dimitrios Papagiannidis, Klára Odehnalová, Natalie Friemel, Bianca M. Esch, Florian Fröhlich, Marcin Luzarowski, Georg H. H. Borner, Friedrich Förster, Sebastian Schuck

**Affiliations:** Heidelberg University Biochemistry Center, Heidelberg, Germany; Present address: Institute of Molecular Systems Medicine, Goethe University Frankfurt, Frankfurt am Main, Germany; Structural Biochemistry, Department of Chemistry, Bijvoet Centre for Biomolecular Research, Utrecht University, Utrecht, Netherlands; Present address: Heidelberg University Biochemistry Center, Heidelberg, Germany; Asklepios BioPharmaceutical, Heidelberg, Germany; Present address: European Molecular Biology Laboratory, Heidelberg, Germany; Department of Biology/Chemistry, Bioanalytical Chemistry Section, Osnabrück University, Osnabrück, Germany; Center for Cellular Nanoanalytic Osnabrück, Osnabrück University, Osnabrück, Germany; Core Facility for Mass Spectrometry and Proteomics, Center for Molecular Biology at Heidelberg University, Heidelberg, Germany; Department of Proteomics and Signal Transduction, Systems Biology of Membrane Trafficking Research Group, Max Planck Institute of Biochemistry, Martinsried, Germany

## Abstract

ESCRT proteins remodel membranes at many cell organelles, including the endoplasmic reticulum (ER). Here, we investigate whether ESCRTs in budding yeast participate in stress-induced ER reorganisation. We find that ER stress triggers the formation of tubular ER subdomains that recruit various ESCRT proteins. Recruitment of the major ESCRT-III protein Snf7 is mediated by the ESCRT-associated protein Bro1, a homologue of human ALIX, in a manner that is mechanistically distinct from Bro1 function at endosomes. ESCRT-containing ER subdomains are derived from ceramide-rich ER exit sites and form contacts with the Golgi. Furthermore, Bro1 helps to concentrate the tethering and lipid transfer protein Tcb3, a homologue of human extended synaptotagmins, at these organelle contacts and contributes to cellular fitness when lipid metabolism is perturbed. These results indicate that specialised ER exit sites can be repurposed for contacting the Golgi directly and uncover ESCRTs as organisers of stress-inducible ER-Golgi contacts that help maintain cell homeostasis.

## Introduction

The endoplasmic reticulum (ER) is a multifunctional organelle with a complex morphology. It consists of flat membrane sheets and high-curvature membrane tubules and contains functionally specialised subdomains, such as sites for protein export and contact with other organelles (Obara et al, 2023). This basic architecture can undergo extensive remodelling. When cells adapt to changing physiological demands, for example during differentiation, ER size can increase several-fold and ER structure can become dominated by sheets or tubules (Fawcett, 1981; Wiest et al, 1990). An important cue for ER remodelling is ER stress resulting from protein misfolding. ER stress activates the unfolded protein response (UPR; Walter and Ron, 2011). UPR activation drives ER membrane expansion, which typically results in the proliferation of sheets (Bernales et al, 2006; Schuck et al, 2009; Ordonez et al, 2013; Wikstrom et al, 2013). The expanded ER can then accommodate new machinery for protein folding and degradation to more effectively eliminate misfolded proteins. Moreover, ER stress can trigger membrane rearrangements that produce new organelle subdomains, including large tubular clusters and multilamellar whorls (Huyer et al, 2004; Schäfer et al, 2020; Xu et al, 2021). Despite numerous studies highlighting the structural plasticity of the ER, the molecular mechanisms of many ER remodelling processes remain unknown.

ESCRTs (endosomal sorting complexes required for transport) constitute an ancient membrane remodelling machinery present in all kingdoms of life (Olmos, 2022; Nachmias et al, 2025; Weiner et al, 2025). ESCRT proteins can associate with various cell membranes, where they assemble into filaments that bring about membrane budding and scission. These activities are essential for a large range of processes, such as endosomal sorting, cytokinesis, autophagy, nuclear envelope reformation, plasma membrane repair and viral egress (Vietri et al, 2020; Hurley et al, 2025). ESCRT proteins usually drive membrane budding away from the cytosol. However, they can also mediate budding towards the cytosol or serve as stable membrane scaffolds (Allison et al, 2013; Penalva et al, 2014; Stempels et al, 2023). At the core of the ESCRT machinery are ESCRT-III proteins, which form heteropolymeric filaments, and the AAA-type ATPase VPS4, which reshapes these filaments (Adell et al, 2017; Mierzwa et al, 2017; Pfitzner et al, 2021). Different pathways exist for membrane recruitment of ESCRT-III. In endosomal sorting, ESCRT-0, -I and -II recognise cargo and act in a recruitment cascade that culminates in membrane association of ESCRT-III. In addition, ESCRT-associated proteins, such as human ALIX and CHMP7, can recruit ESCRT-III, either alone or in collaboration with ESCRT-0, -I and -II (Schöneberg et al, 2017).

The ESCRT machinery has diverse roles at the ER. For instance, ESCRT-III proteins in budding yeast help release pre-peroxisomal vesicles from the ER (Mast et al, 2018). In mammalian cells under nutrient stress, ESCRT machinery appears to associate with COPII coat proteins to mediate selective microautophagy of ER exit sites (Liao et al, 2024). ESCRT proteins have also been linked to COPII function and autophagy in flies (Wang et al, 2022). Finally, viruses manipulate the ESCRT machinery of their hosts to generate viral replication compartments at the ER and bud into the ER lumen (Diaz et al, 2015; Tabata et al, 2016).

Here, we investigate whether ESCRT proteins are involved in ER remodelling during ER stress in budding yeast. We provide evidence that ESCRT proteins function as organisers of stress-induced ER-Golgi contacts and suggest that these contacts promote non-vesicular lipid export from the ER to maintain organelle homeostasis.

## Results

### The ESCRT-III protein Snf7 is recruited to stress-induced ER subdomains

To explore a possible role of the ESCRT machinery in stress-induced ER remodelling, we examined Snf7, an abundant ESCRT-III protein essential for all known ESCRT functions (Teis et al, 2008; Heinzle et al, 2019; Platzek et al, 2025; Hurley et al, 2025). Fusing Snf7 with a fluorescent protein renders it inactive (Teis et al, 2008). However, tagged Snf7 molecules can serve as tracers for functional ESCRT-III assemblies if they are present alongside endogenous Snf7 molecules without outnumbering them (Adell et al, 2017). Accordingly, we fused Snf7 with the fluorescent proteins mNeonGreen or mScarlet-i (referred to as neon or scarlet) and expressed it under the *VPS24* promoter in cells also containing endogenous Snf7. The levels of tagged Snf7 were substantially lower than those of endogenous Snf7 (Figure S1A). We showed previously that this amount of Snf7-neon did not impair endosomal sorting or micro-ER-phagy (Schäfer et al, 2020), and we confirmed here that also Snf7-scarlet did not disturb endosomal sorting (Figure S1B).

Snf7-neon localised to the cytosol and endosomes, as reported (Figure S1C; Adell et al, 2017). We then treated cells with tunicamycin to block protein N-glycosylation, cause secretory protein misfolding and thus induce ER stress. Strikingly, Snf7-neon formed puncta substantially brighter than the endosomal Snf7 signal. These bright Snf7 puncta consistently co-localised with regions densely labelled by the ER marker cherry-Ubc6, either at the nuclear envelope or the peripheral ER (Figure 1A). We refer to these regions as ER clusters. Snf7 occasionally also formed dim puncta at the nuclear envelope, which may reflect the role of ESCRT proteins in nuclear pore complex quality control (Figure S1D; Webster et al, 2016). However, these structures did not co-localise with ER clusters and we did not investigate them further. Importantly, ER recruitment of Snf7-neon to ER clusters was lost in the absence of endogenous Snf7 and hence involved the activity of native Snf7 (Figure 1B). This observation argues against the possibility that the formation of bright Snf7-neon puncta simply reflected aggregation of non-functional Snf7.

**Figure 1.**
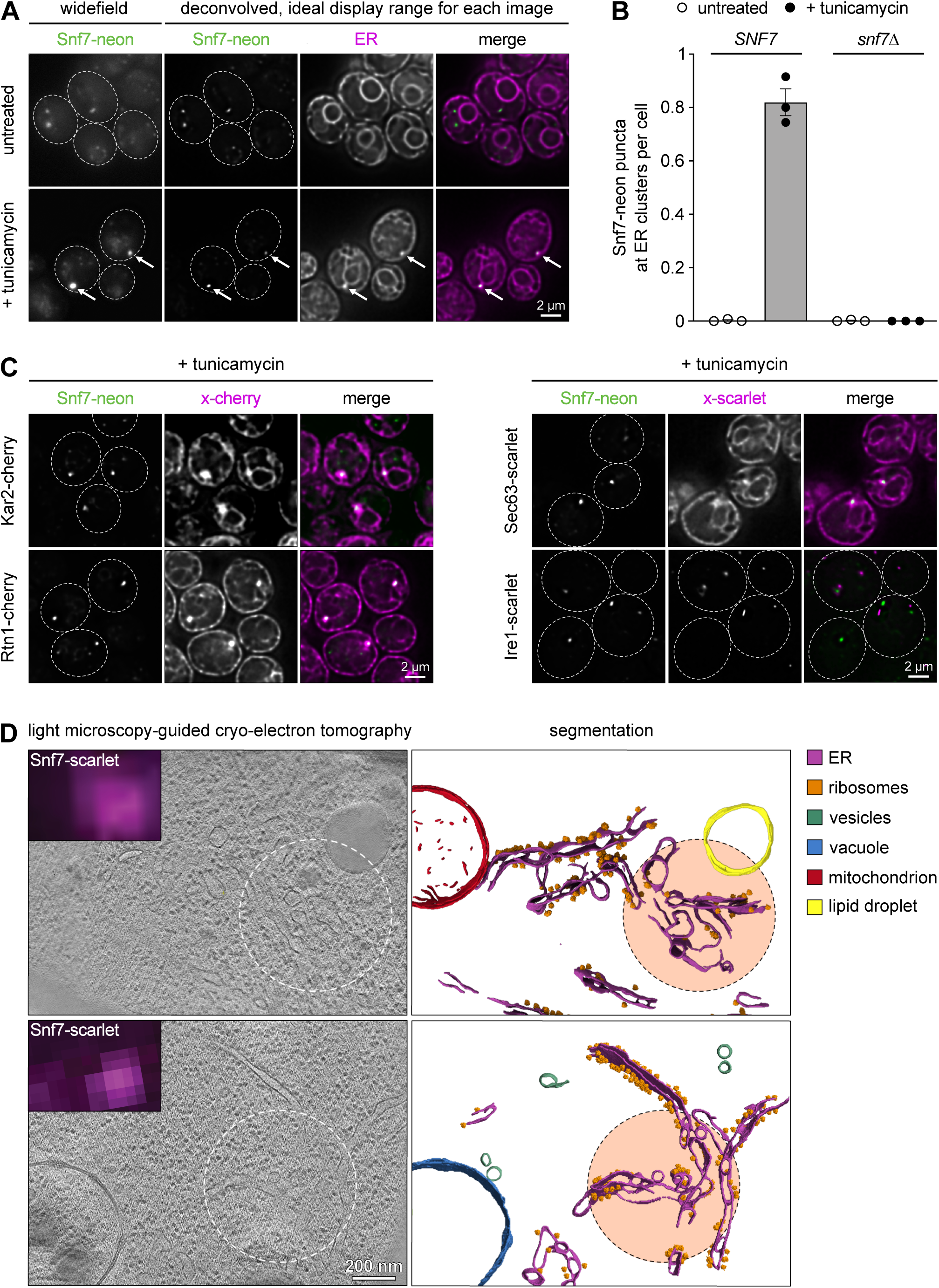
The ESCRT-III protein Snf7 is recruited to stress-induced ER subdomains. **(A)** Widefield fluorescent images and deconvolved fluorescent images of untreated and tunicamycin-treated wild-type cells expressing Snf7-neon under the control of the *VPS24* promoter along with the general ER marker cherry-Ubc6. Widefield images were adjusted to the same display range for untreated and tunicamycin-treated cells to highlight the brightness difference between endosomal and ER-localised Snf7 puncta. Deconvolved images were adjusted to the ideal display range for each image. Thus, intensities cannot be compared across images. Dotted lines indicate cell boundaries. Arrows mark Snf7-neon puncta at ER clusters. **(B)** Quantification of Snf7-neon puncta at ER clusters per cell in untreated and tunicamycin-treated cells expressing Snf7-neon under the control of the *VPS24* promoter in presence (*SNF7*) or absence (s*nf7Δ*) the endogenous *SNF7* gene. Data are the mean of n = 3 biological replicates, error bars indicate the standard error of the mean. **(C)** Deconvolved fluorescent images of tunicamycin-treated cells expressing Snf7-neon and the ER proteins Kar2, Rtn1, Sec63 or Ire1 fused to the red fluorescent proteins cherry or scarlet. Dotted lines indicate cell boundaries. **(D)** Widefield fluorescent images (inset top left), tomographic slices corresponding to the same field of view (left) and segmentation with ribosome mapping (right) of tunicamycin-treated cells expressing Snf7-scarlet. Dotted lines indicate the centre of the Snf7-scarlet fluorescent signal.

Overall Snf7 protein levels did not change during stress (Figure S1A), indicating that the bright Snf7 puncta arose from redistribution of Snf7 to the ER. Puncta formed within one hour of tunicamycin treatment and, after three hours, the majority of cells had a single Snf7 punctum at an ER cluster (Figure S1E). Cells with two or more Snf7 puncta were rare (Figure S1F). Dithiothreitol treatment, which causes ER stress by preventing disulfide bond formation, induced very little Snf7 recruitment to the ER (Figure S1E), and we therefore used tunicamycin in subsequent experiments.

Stress-induced Snf7 puncta were positive for the lumenal ER chaperone Kar2, the ER-tubulating reticulon protein Rtn1 and the ER transmembrane protein Sec63 (Figure 1C). By contrast, they showed no overlap with clusters of the ER stress sensor Ire1 or the autophagosome marker Atg8 (Figures 1C and S1G). In addition, Snf7 puncta still formed when macroautophagy was disabled by removal of Atg7 (Figure S1H). Hence, Snf7 puncta formation and macroautophagy are separate processes. To determine ER ultrastructure at sites of Snf7 recruitment, we applied light microscopy-guided cryo-electron tomography. Reconstructions revealed dense, irregularly shaped, mostly tubular membrane structures (Figure 1D). These structures were bound by ribosomes, which identified them as ER. Interestingly, the fluorescent Snf7 signal best correlated with ER subregions partially devoid of ribosomes, implying structural and functional specialisation. None of the Snf7 puncta corresponded to multilamellar ER whorls, indicating that they were not linked to microautophagy of the ER (Schäfer et al, 2020).

In summary, ER stress triggers recruitment of the ESCRT-III protein Snf7 to tubular ER subdomains that appear unrelated to Ire1-mediated stress sensing or autophagy.

### ER recruitment of Snf7 requires the ESCRT-associated proteins Bro1 and Rim20

Next, we investigated the mechanism of stress-induced recruitment of Snf7 to the ER. The process did not depend on ESCRT-0, -I, -II or the ESCRT-III protein Vps20, which links ESCRT-II and -III (Hurley et al, 2025). The number of ER-localised Snf7 puncta even increased in the absence of these ESCRTs, presumably because Snf7 was released from endosomes and more Snf7 was available for recruitment to the ER. Chm7, which brings ESCRT-III to the nuclear envelope, was also not needed for Snf7 recruitment. By contrast, the ALIX family protein Bro1 was strictly required (Figure 2A). ER clusters persisted in *bro1Δ* cells, showing that they can form independently of Snf7 recruitment (Figures 2B and S2A). Bro1 contains a Bro1 domain, which it shares with two other yeast proteins, Rim20 and Ygr122w (Kim et al, 2005; Galindo et al, 2007). The number of Snf7 puncta at ER clusters was unaffected by removal of Ygr122w but strongly reduced by removal of Rim20 (Figure 2A). Membrane recruitment of ESCRT-III by Bro1 domain proteins independently of ESCRT-0/I/II is known in mammalian cells (Schöneberg et al, 2017). However, stress-induced Snf7 recruitment to the ER is the first such process uncovered in yeast.

**Figure 2.**
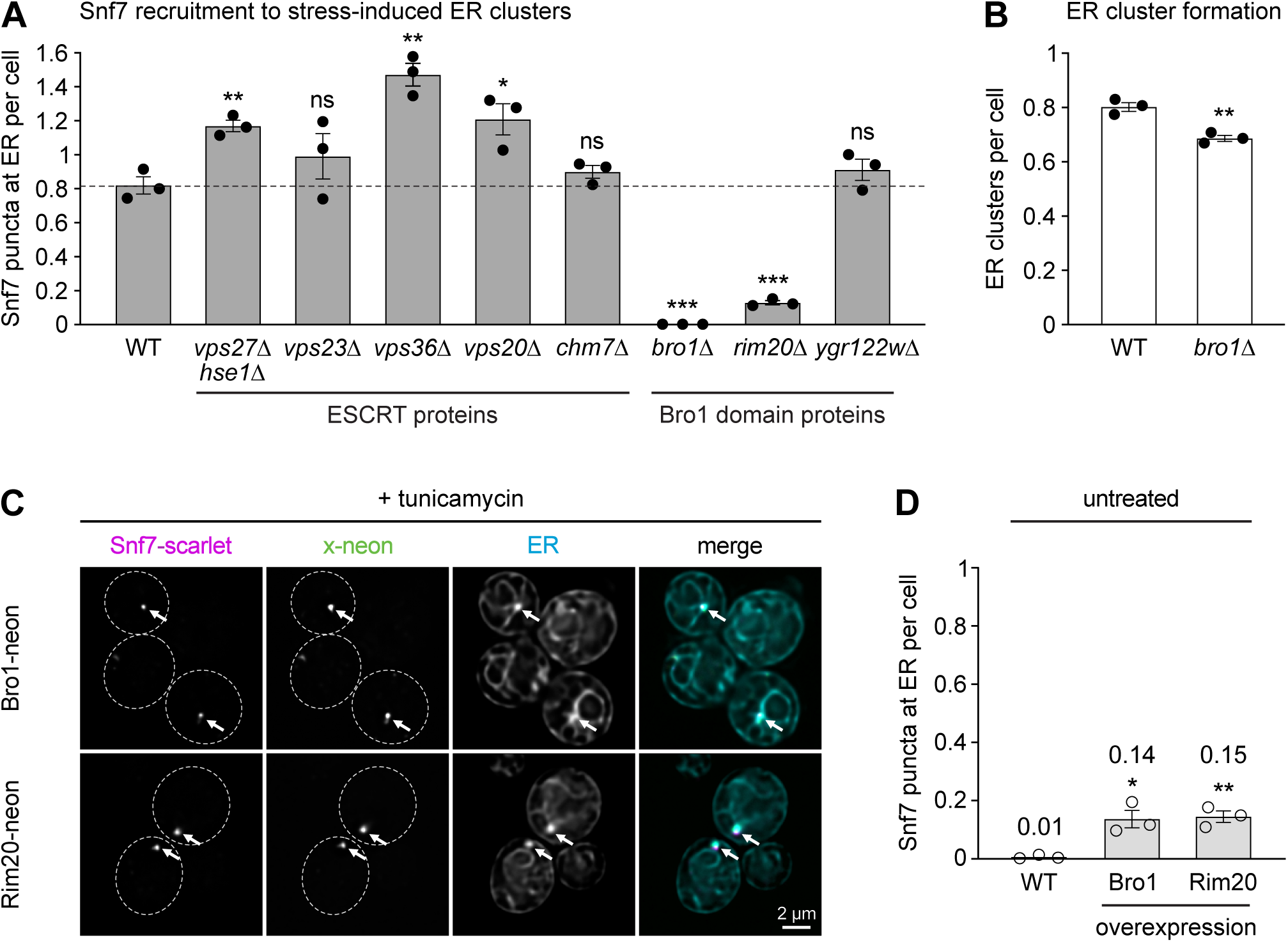
ER recruitment of Snf7 requires the ESCRT-associated proteins Bro1 and Rim20. **(A)** Quantification of Snf7-neon puncta at ER clusters per cell in tunicamycin-treated wild-type (WT) cells and mutants lacking the indicated ESCRT or Bro1 domain proteins. Vps27 and Hse1 are ESCRT-0 proteins, Vps23 is an ESCRT-I protein, Vps36 is an ESCRT-II protein, Vps20 links ESCRT-II and ESCRT-III, and Chm7 is an ESCRT-II/III chimera (Webster et al, 2016). Data are the mean of n = 3 biological replicates, error bars indicate the standard error of the mean. For each mutant, statistical significance of the difference to wild-type cells was evaluated with a two-tailed homoscedastic t-test. * p<0.05, ** p<0.01, *** p<0.001, ns = not significant. **(B)** Quantification of ER clusters per cell in WT and *bro1Δ* cells. Data are the mean of n = 3 biological replicates, error bars indicate the standard error of the mean. Statistical significance of the difference between wild-type and *bro1Δ* cells was evaluated with a two-tailed homoscedastic t-test. **(C)** Deconvolved fluorescent images of tunicamycin-treated cells expressing Snf7-scarlet, the ER marker BFP-Ubc6 and either Bro1-neon or Rim20-neon. Dotted lines indicate cell boundaries. Arrows mark Snf7-scarlet puncta. **(D)** Quantification of Snf7-neon puncta at ER clusters per cell in untreated wild-type, Bro1-overexpressing and Rim20-overexpressing cells. Data are the mean of n = 3 biological replicates, error bars indicate the standard error of the mean. Statistical significance of the difference between the wild-type and each overexpression strain was evaluated with a two-tailed homoscedastic t-test.

Bro1 and Rim20 both formed stress-induced puncta, which co-localised with Snf7 at ER clusters (Figure 2C). These puncta formed also in the absence of tagged Snf7, albeit in a much smaller fraction of cells (Figure S2B, C). Hence, ER recruitment of Bro1 and Rim20 can occur in cells possessing only endogenous Snf7, but tagged Snf7 appears to stabilise their association with the ER. Membrane recruitment of ESCRT proteins is often highly dynamic and transient (Adell et al, 2017), and the same may be true for ER recruitment of Bro1 and Rim20 in the absence of tagged Snf7. Both Bro1 and Rim20 interact with Snf7 via their Bro1 domains (Xu et al, 2004; Kim et al, 2005). In addition, Bro1 and Rim20 puncta at the ER frequently overlapped (Figure S2D). We therefore tested whether the two proteins had related functions in Snf7 recruitment to the ER. Bro1 overexpression in *rim20Δ* cells and Rim20 overexpression in *bro1Δ* cells restored some Snf7 puncta formation. This result indicated that Bro1 and Rim20 can partially compensate for each other in recruiting Snf7 to ER subdomains (Figure S2E). Finally, both Bro1 and Rim20 overexpression induced Snf7-containing ER clusters in about 15% of cells in the absence of ER stress (Figures 2D and S2F), emphasising the key role of Bro1 domain proteins in Snf7 recruitment.

The data so far showed that Bro1 and Rim20 both have roles in ER recruitment of Snf7 but that Bro1 is strictly necessary. Bro1 additionally acts in endosomal sorting (Odorizzi et al, 2003). To uncouple these two functions, we searched for features of Bro1 that are specifically needed to bring Snf7 to the ER. Bro1 contains an N-terminal Bro1 domain, a V domain and a proline-rich region (Figure 3A). The V domain and proline-rich region were not essential for Snf7 recruitment (Figure S3). The Bro1 domain contains a tetratricopeptide repeat (TPR)-like domain and two hydrophobic surface patches. The first patch physically interacts with Snf7, the second has no known function (Figure 3A; Kim et al, 2005). The TPR-like domain and the second hydrophobic patch can be disrupted by K246A and Y320D mutations, respectively (Kim et al, 2005), and we individually introduced these mutations into the endogenous *BRO1* gene. Both bro1(K246A) and bro1(Y320D) were fully functional in endosomal sorting but could not recruit Snf7 to the ER during stress (Figure 3B, C). Moreover, they failed to localise to the ER, even in cells expressing tagged Snf7 (Figure 3D). Thus, the TPR-like domain and the second hydrophobic patch are dispensable for endosomal sorting but required for stress-induced ER association of Bro1.

**Figure 3.**
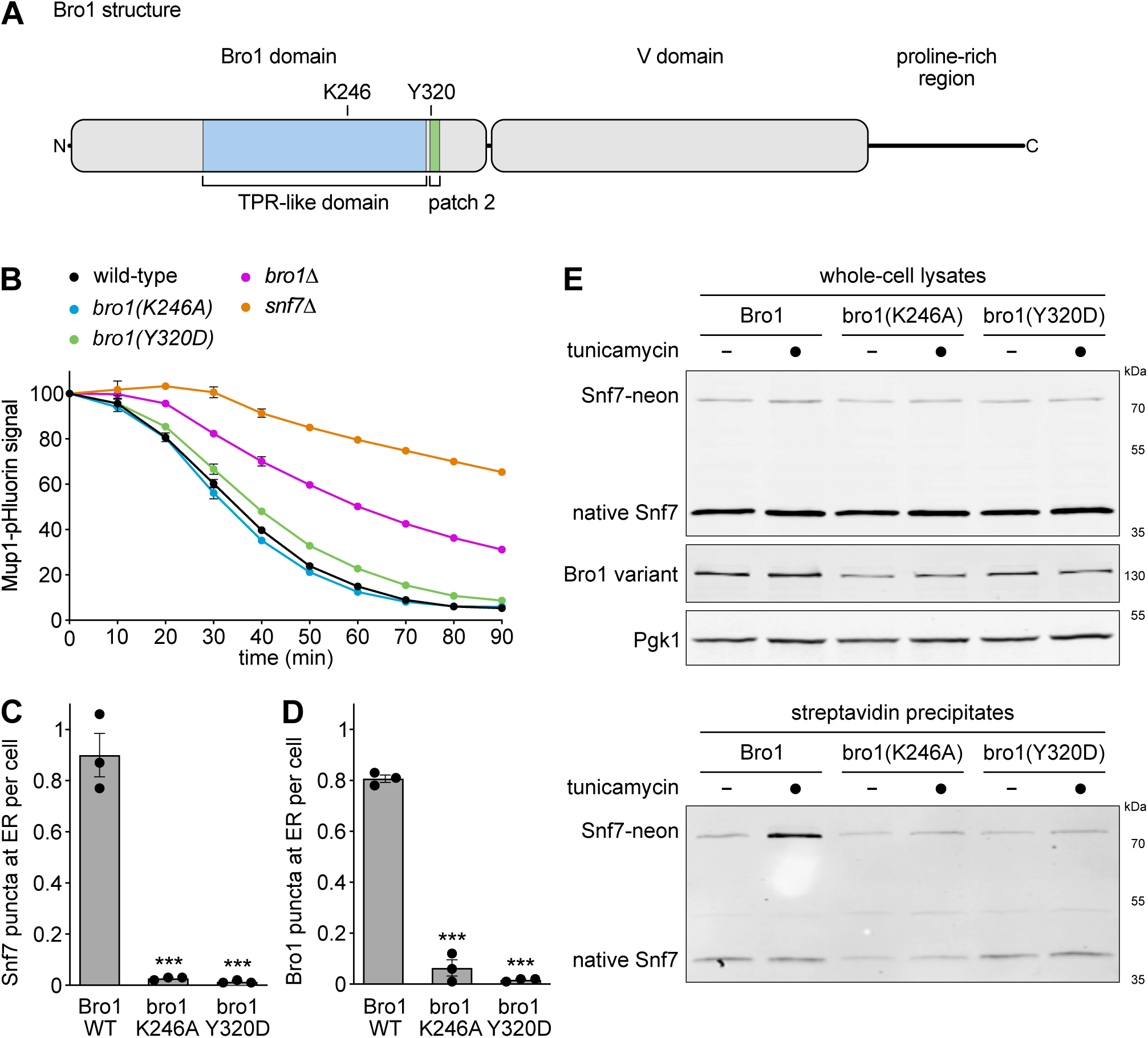
Snf7 recruitment to the ER requires specific features of Bro1. **(A)** Schematic of Bro1 structure. TPR, tetratricopeptide repeat. Patch 2 refers to a hydrophobic surface with unknown function. The residues constituting hydrophobic patch 1 are non-contiguous and are distributed between residues 128 and 351. **(B)** Endosomal sorting assay based on Mup1-pHluorin fluorescence, as measured by flow cytometry. The x-axis indicates the time after addition of methionine to induce ESCRT-dependent transport of Mup1-pHluorin to the vacuole, where the pHluorin signal is quenched. For each strain, fluorescence was normalised to t = 0. Data are the mean of n = 3 biological replicates, error bars indicate the standard error of the mean. **(C)** Quantification of Snf7-neon puncta at ER clusters in tunicamycin-treated wild-type (WT) cells and cells expressing the indicated mutant versions of Bro1. Data are the mean of n = 3 biological replicates, error bars indicate the standard error of the mean. Statistical significance of the difference between cells expressing wild-type Bro1 and each Bro1 variant was evaluated with a two-tailed homoscedastic t-test. *** p<0.001. **(D)** As in panel C but quantification of Bro1-neon puncta at ER clusters. **(E)** Western blots of Snf7, myc tag and Pgk1 from a Snf7 biotinylation assay. Cells expressing Snf7-neon and wild-type Bro1, bro1(K246A) or bro1(Y320D) fused to TurboID-myc were treated with tunicamycin where indicated. Whole cell lysates were analysed to determine expression levels. Pgk1 served as a loading control. Biotinylated proteins were precipitated with streptavidin beads and analysed to determine the extent of Snf7 biotinylation and hence Bro1-Snf7 proximity.

To establish a biochemical readout for Bro1-Snf7 proximity, we fused wild-type Bro1, bro1(K246A) and bro1(Y320D) to the biotin ligase TurboID and assayed their ability to biotinylate endogenous Snf7 and Snf7-neon. At steady state, the three Bro1 variants biotinylated Snf7 equally, likely reflecting proximity of Bro1 and Snf7 at endosomes. ER stress increased biotinylation of Snf7-neon by wild-type Bro1, presumably as a result of their accumulation at ER clusters. Only a marginal increase of biotinylation was observed with the two mutant Bro1 variants (Figure 3E). These data support the notion that bro1(K246A) and bro1(Y320D) were specifically defective in functioning at the ER. Biotinylation of endogenous Snf7 remained unchanged upon stress, possibly because tagged Snf7 was enriched over endogenous Snf7 at ER clusters.

These findings established that stress-induced recruitment of the ESCRT-III protein Snf7 to ER subdomains is independent of ESCRT-0/I/II, requires ESCRT-associated Bro1 and Rim20 in a partially overlapping manner, and depends on the TPR-like domain and the second hydrophobic patch within the Bro1 domain of Bro1.

### Constituents of Snf7-containing ER subdomains and factors needed for their formation

To identify additional constituents of Snf7-containing ER subdomains, we carried out a proximity-dependent biotinylation screen. We tagged Bro1 or recruitment-deficient bro1(K246A) with TurboID, expressed Snf7-neon, and blocked association of Bro1 with endosomes or sites of nuclear pore complex quality control by removing Vps27 and Chm7 (Katzmann et al, 2003; Webster et al, 2016). Furthermore, we eliminated the vacuolar proteases Pep4 and Prb1 to reduce proteolysis in living cells or after cell lysis. Cells were left untreated or treated with tunicamycin, and biotinylated proteins were identified by mass spectrometry (Figure 4A). 155 proteins were biotinylated upon ER stress by Bro1 but not bro1(K246A). Among these hits were Bro1 itself, Snf7, Rim20 and four other ESCRT-III or ESCRT-associated proteins (Figure 4B; Table S1). These data implied that Snf7 was recruited to the ER as part of ESCRT-III assemblies. Accordingly, the ESCRT-III protein Ist1, which can be tagged without disrupting its localisation (Dimaano et al, 2008), partially co-localised with Snf7 at ER clusters (Figure 4C). In addition, hits included five COPI proteins for transport within the Golgi and between Golgi and ER, two COPII proteins for ER-to-Golgi transport, five ER proteins including Rtn1 and Sec63, and six Golgi proteins (Figure 4B; Table S1).

**Figure 4.**
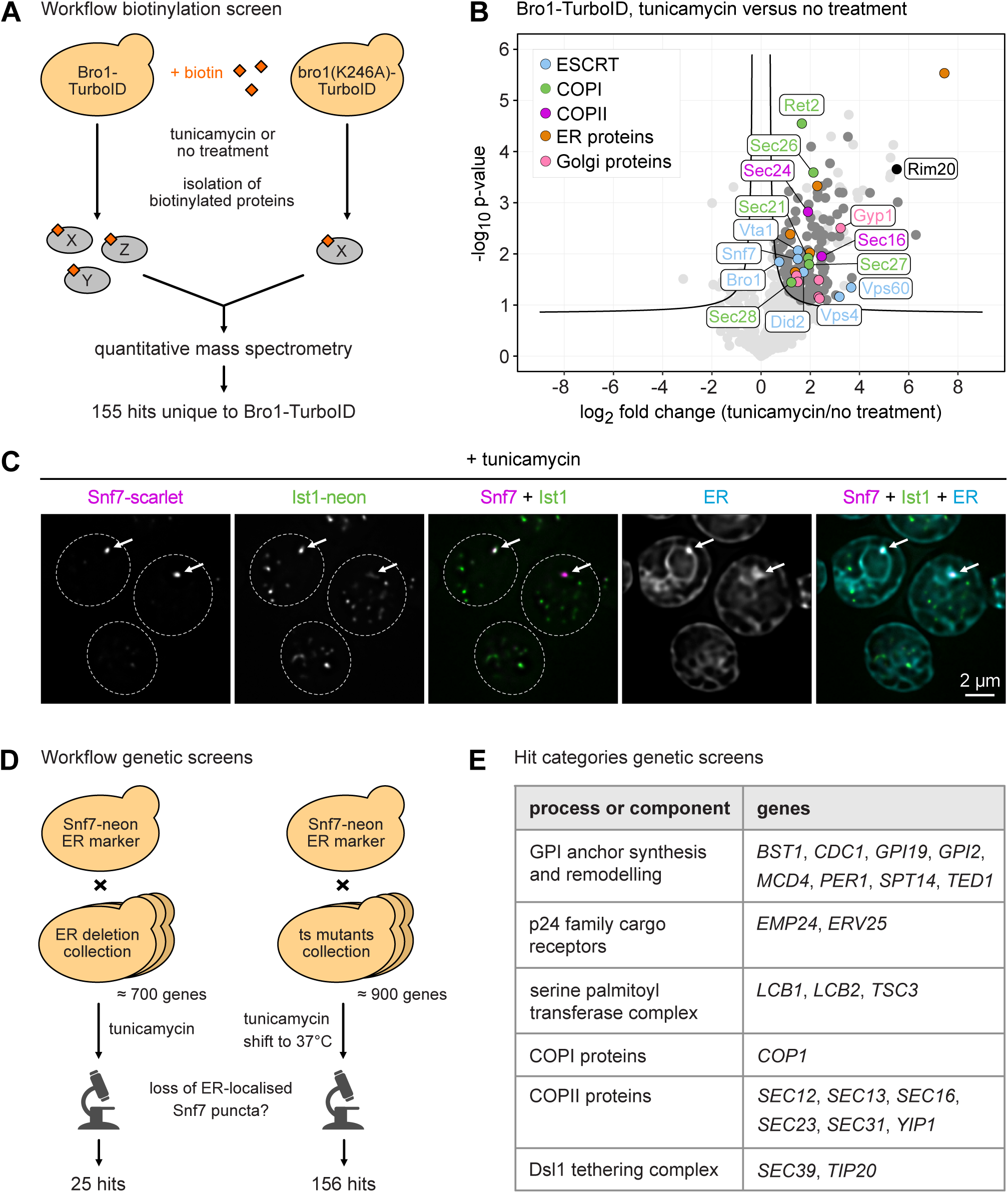
Constituents of Snf7-containing ER subdomains and factors needed for their formation. **(A)** Workflow of the biotinylation screen. **(B)** Volcano plot of biotinylated proteins retrieved from Bro1-TurboID cells, showing log_2_ fold changes upon tunicamycin treatment. Volcano lines indicate 5% false discovery rate cut-offs based on data permutation. Hits specific to Bro1-TurboID are indicated in dark grey or in colour. Non-hits and hits shared between Bro1-TurboID and bro1(K246A)-TurboID are indicated in light grey. **(C)** Deconvolved fluorescent images of tunicamycin-treated cells expressing Snf7-scarlet, Ist1-neon and the ER marker BFP-Ubc6. Dotted lines indicate cell boundaries. Arrows mark Snf7-scarlet puncta. **(D)** Workflow of the genetic screens with the collections of ER-related gene deletion mutants (ER deletion collection) and temperature-sensitive mutants (ts mutants collection). **(E)** Hit categories from the genetic screens.

To identify factors needed to generate Snf7-containing ER subdomains, we carried out two genetic screens. We introduced Snf7-neon and cherry-Ubc6 into collections of about 700 mutants lacking non-essential genes related to ER function (Table S2A) and about 900 temperature-sensitive mutants of essential and non-essential genes (Li et al, 2011; Table S3A). Using automated microscopy, we searched for mutants with defects in the formation of tunicamycin-induced Snf7 puncta at ER clusters (Figure 4D). Screening of the ER-related deletion mutants yielded 25 hits, including three genes encoding proteins for glycosylphosphatidylinositol (GPI) anchor remodelling and two genes encoding p24 family cargo receptors for ER export of GPI-anchored proteins (Table S2B). Screening of the temperature-sensitive mutants yielded 156 hits. Exclusion of genes with general functions in chromatin remodelling, DNA replication, chromosome segregation, transcription or translation reduced this number to 68 hits. Among these were genes encoding five proteins for GPI anchor synthesis, three subunits of the serine palmitoyltransferase complex for sphingolipid synthesis, one COPI protein, six proteins related to COPII vesicles and two subunits of the Dsl1 tethering complex for Golgi-to-ER transport (Table S3B). Taken together, the two genetic screens identified 22 genes in the hit categories listed above (Figure 4E).

Overall, the biotinylation and genetic screens suggested that the formation of ESCRT-III assemblies at the ER is linked, both physically and functionally, to a large number of proteins acting at the interface of ER and Golgi.

### Cooperation of proteins at the ER-Golgi interface in ESCRT-III recruitment to the ER

The genetic screens identified several factors involved in ER export of GPI-anchored proteins (GPI-APs). The sorting and transport of GPI-APs involves a lipid-based mechanism. When a GPI anchor has been attached to a protein, its lipid and glycan parts are modified by GPI anchor remodellers. The finished GPI anchor then serves as an ER exit signal. Since GPI-APs have no cytosolic domain, they cannot interact with the general ER export machinery directly. However, GPI-APs associate with long-chain ceramides and partition into ordered membrane microdomains that function as specialised ER exit sites. There, GPI-APs bind to cargo receptors of the p24 family, which form heteromeric complexes, span the ER membrane and interact with COPII proteins, thus linking GPI-APs to the COPII coat. As a result, special COPII transport carriers are generated that are enriched in ceramide and GPI-APs (Lopez et al, 2019; Rodriguez-Gallardo et al, 2020).

In agreement with the genetic screens, removal of the GPI anchor remodellers Ted1 or Bst1 reduced the number of stress-induced Snf7 puncta at the ER (Figure 5A). Removal of the screen hit Tsc3, an auxiliary subunit of the serine palmitoyltransferase complex needed for efficient synthesis of sphingolipids (Gable et al, 2000), including ceramide, yielded an even stronger phenotype. Snf7 puncta were also less numerous in mutants lacking the p24 cargo receptors Emp24 or Erv25. Furthermore, all mutants had fewer ER clusters than wild-type cells (Figure 5B). This finding implied that stress-induced formation of ER clusters involved membrane microdomains comprising ceramide, GPI-APs and p24 proteins. If biogenesis of these domains was disturbed, ER cluster formation and Snf7 recruitment to ER clusters were diminished.

**Figure 5.**
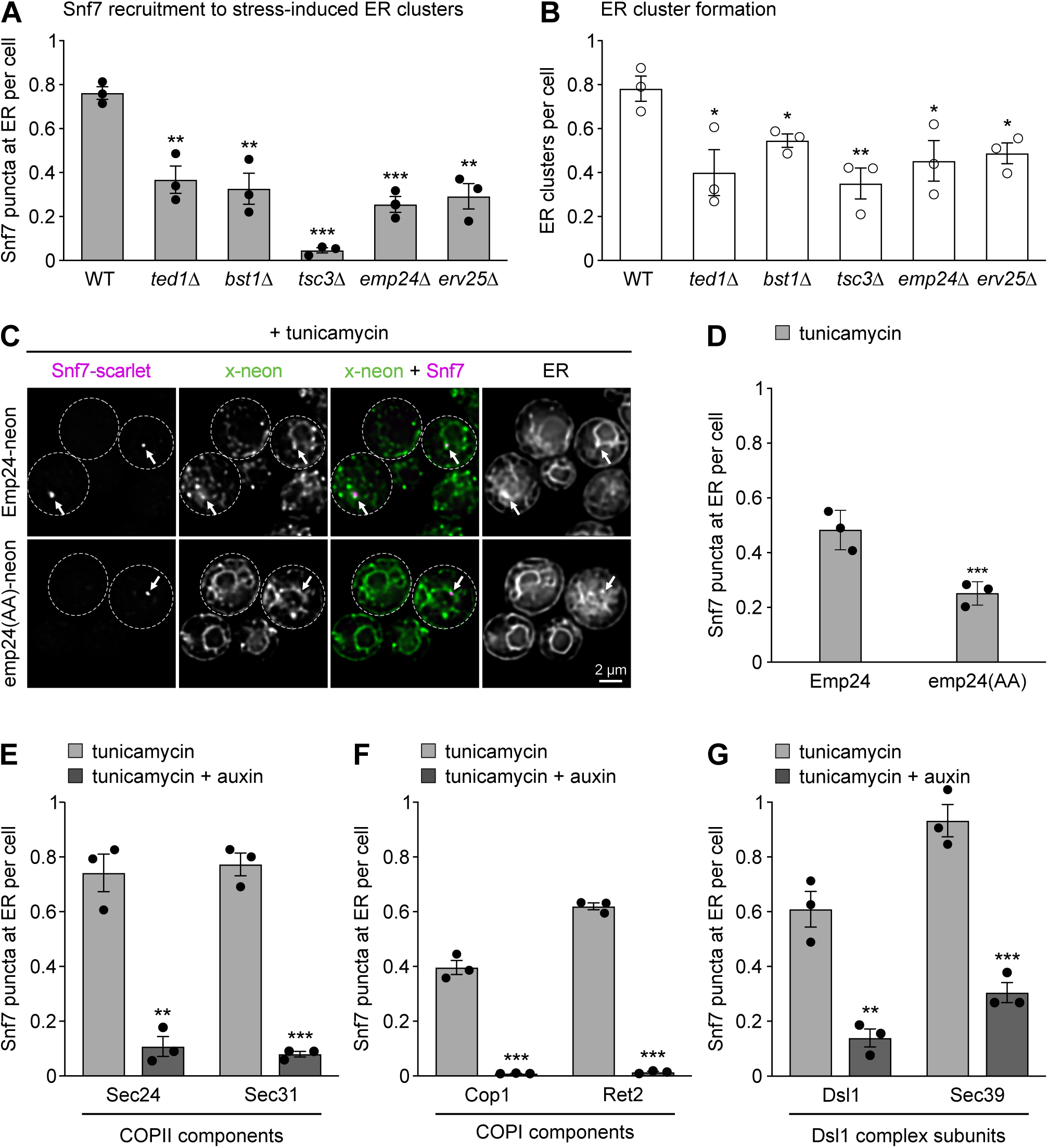
Machinery for transport between ER and Golgi is required for ER recruitment of Snf7. **(A)** Quantification of Snf7-neon puncta at ER clusters in tunicamycin-treated wild-type (WT) cells and the indicated mutants. Data are the mean of n = 3 biological replicates, error bars indicate the standard error of the mean. For each mutant, statistical significance of the difference to wild-type cells was evaluated with a two-tailed homoscedastic t-test. * p<0.05, ** p<0.01, *** p<0.001. **(B)** As in panel A but quantification of ER clusters marked with cherry-Ubc6. **(C)** Deconvolved fluorescent images of cells expressing Snf7-scarlet, the ER marker BFP-Ubc6 and either Emp24-neon or emp24(AA)-neon. Dotted lines indicate cell boundaries. Arrows mark Snf7-scarlet puncta. **(D)** Quantification of Snf7-scarlet puncta at ER clusters in tunicamycin-treated Emp24-neon and emp24(AA)-neon cells. Data are the mean of n = 3 biological replicates, error bars indicate the standard error of the mean. Statistical significance of the difference between cells expressing Emp24-neon and emp24(AA)-neon was evaluated with a two-tailed homoscedastic t-test. Snf7 puncta are less frequent already in Emp24-neon cells (compare with WT cells in panel A), likely reflecting that tagged Emp24 is only partially functional. **(E)** Quantification of Snf7-neon puncta at ER clusters in tunicamycin-treated cells without and with auxin-induced degradation of the COPII proteins Sec24 or Sec31. Data are the mean of n = 3 biological replicates, error bars indicate the standard error of the mean. Statistical significance of the difference between cells with and without auxin treatment was evaluated with a two-tailed homoscedastic t-test. **(F)** As in panel E but for auxin-induced degradation of the COPI proteins Cop1 and Ret2. Snf7 puncta are less frequent already in strains without Cop1 and Ret2 depletion, likely reflecting that degron-tagged Cop1 and Ret2 are only partially functional. **(G)** As in panel E but for the Dsl1 complex subunits Dsl1 and Sec39. Snf7 puncta are less frequent already in cells without Dsl1 depletion, likely reflecting that degron-tagged Dsl1 is only partially functional.

The cytosolic C-terminus of Emp24 contains a diaromatic motif that binds to the COPII coat (Belden and Barlowe, 2001). To determine whether this interaction was relevant for ER association of Snf7, we asked whether Emp24-neon and emp24(AA)-neon, which lacks the diaromatic motif, co-localised with Snf7-scarlet at ER clusters. Emp24-neon was enriched at stress-induced Snf7-positive ER clusters (Figure 5C). By contrast, emp24(AA)-neon localised throughout the ER, without enrichment at Snf7-containing subdomains. Furthermore, emp24(AA)-neon cells showed fewer Snf7 puncta (Figure 5C, D). Hence, interaction of p24 family cargo receptors with the COPII coat was important for ER recruitment of Snf7.

Next, we tested the requirement for COPII coat components. Since COPII proteins are encoded by essential genes, we targeted Sec24 and Sec31 by means of an auxin-inducible protein degradation system (Figure S4A; Hubbe et al, 2026). Induced degradation of Sec24 and Sec31 strongly reduced the number of Snf7 puncta at the ER (Figure 5E). Overall ER morphology was severely disturbed by Sec24 and Sec31 depletion, so we could not determine whether there was a specific effect on stress-induced formation of ER clusters (Figure S4B).

Two further groups of screen hits participate in a shared pathway: COPI proteins and subunits of the Dsl1 tethering complex. In retrograde Golgi-ER transport, COPI vesicles form at the Golgi, are captured by the Dsl1 complex at ER arrival sites and then fuse with the ER (Spang, 2009; Schröter et al, 2016). Auxin-induced depletion of the essential COPI proteins Cop1 and Ret2 drastically reduced recruitment of Snf7 to the ER (Figures 5F and S4A). Similarly, depletion of two essential subunits of the Dsl1 complex, Dsl1 and Sec39, strongly reduced the number of Snf7 puncta at the ER (Figures 5G and S4A). Depletion of both COPI proteins and Dsl1 complex subunits generally disturbed ER morphology, again preventing a quantification of ER clusters.

In summary, stress-induced formation of ER subdomains and recruitment of ESCRT proteins to these sites requires GPI anchor remodellers, proper sphingolipid synthesis, p24 cargo receptors, COPII proteins, COPI proteins and the Dsl1 tethering complex. The requirement for these factors, the importance of the Emp24-COPII interaction for Snf7 recruitment and the co-localisation of Snf7 with Emp24 indicate that Snf7 is recruited to specialised ceramide-rich ER exit sites for GPI-anchored proteins.

### ER subdomains containing Snf7 are part of stress-induced ER-Golgi contacts

Next, we investigated the spatial relationship of factors involved in the generation of Snf7-containing ER subdomains. We first visualised Snf7 and the ER together with COPII and COPI proteins. The COPII protein Sec24-HaloTag localised to small puncta, which frequently abutted or surrounded Snf7-scarlet puncta at ER clusters. In the same cells, the COPI protein Cop1-neon often showed overlap with Snf7-scarlet at ER clusters (Figures 6A and S5). Dsl1-neon was also present at ER clusters containing Snf7-scarlet (Figure 6B).

**Figure 6.**
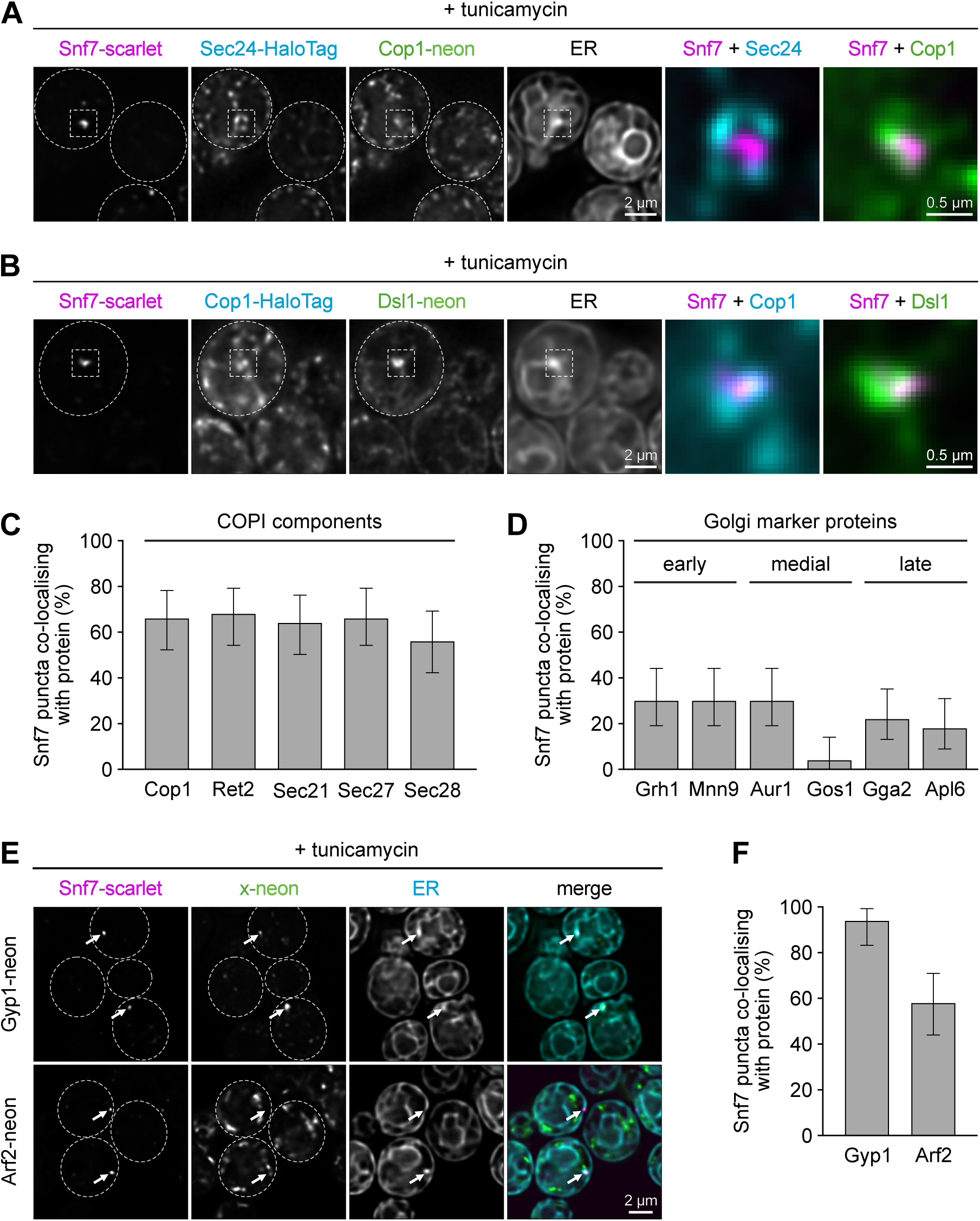
Snf7-containing ER subdomains are part of stress-induced ER-Golgi contacts. **(A)** Deconvolved fluorescent images of tunicamycin-treated cells expressing Snf7-scarlet, the COPII protein Sec24-HaloTag, the COPI protein Cop1-neon and the ER marker BFP-Ubc6. The last two images show magnifications of the area marked with grey boxes in the first four images. **(B)** As in panel A but of cells expressing Snf7-scarlet, Cop1-HaloTag, the Dsl1 complex subunit Dsl1-neon and BFP-Ubc6. **(C)** Quantification of co-localisation of Snf7-scarlet and different COPI proteins in tunicamycin-treated cells. Error bars indicate the margin of error at the 95% confidence level from n = 1 experiment and 50 Snf7 puncta assessed for co-localisation with each COPI protein. **(D)** As in panel C but for co-localisation of Snf7-scarlet and marker proteins of the early, medial and late Golgi. **(E)** Deconvolved fluorescent images of tunicamycin-treated cells expressing Snf7-scarlet, BFP-Ubc6 and either Gyp1-neon or Arf2-neon. Dotted lines indicate cell boundaries. Arrows mark Snf7-scarlet puncta. **(F)** Quantification of co-localisation of Snf7-scarlet and the Golgi proteins Gyp1-neon and Arf2-neon. Error bars indicate the margin of error at the 95% confidence level from n = 1 experiment and 50 Snf7 puncta assessed for co-localisation with each Golgi protein.

Proximity of COPII proteins, COPI proteins and the Dsl1 complex has been observed in yeast before (Schröter et al, 2016; Roy Chowdhury et al, 2020). These data have led to the idea that COPII-positive ER exit sites and COPI-positive ER arrival sites form bidirectional transport portals at closely apposed ER and Golgi membranes (Roy Chowdhury et al, 2020). Therefore, we asked whether co-localisation of stress-induced ER subdomains with COPI proteins reflected proximity of ER arrival sites and free COPI vesicles, or instead indicated proximity of ER and COPI-positive Golgi membranes. Quantification revealed that about 60% of Snf7 puncta at the ER contained COPI proteins, consistent with both possibilities (Figure 6C). Furthermore, Snf7 co-localised partially with marker proteins of the early, medial and late Golgi (Figure 6D). Gyp1, which is found throughout the maturing Golgi and was the strongest Golgi hit from the biotinylation screen, was present at nearly all Snf7 puncta (Figure 6E, F; Thomas et al, 2021; Figure 4B and Table S1). Gyp1 is recruited to the Golgi by the small GTPase Arf1 (Thomas et al, 2021). Arf1 was difficult to visualise because its tagging disturbed ER and Golgi structure. However, the Arf1 paralog Arf2 frequently co-localised with Snf7 puncta at the ER (Figure 6E, F). These results suggested that Snf7 is recruited to sites of close apposition of ER and various Golgi compartments. Sites of convergence of COPI and COPII machinery can act as transport hubs (Kurokawa et al, 2014; Roy Chowdhury et al, 2020). We therefore asked whether Snf7-containing ER subdomains functioned in ER-Golgi transport, in particular of GPI-APs. We imaged ER export of the model GPI-AP Gas1 but found no evidence for a role of Bro1 in the process. We then considered the possibility that the structures we had described as ER clusters were parts of ER-Golgi contacts. It has been shown that ER stress induces ER-Golgi contacts containing the tethering proteins Nvj2 and Tcb3 (Liu et al, 2017, Ikeda et al, 2020). Both Nvj2 and Tcb3 are integral ER membrane proteins. Nvj2 localises to the general ER and nucleus-vacuole junctions at steady state but re-localises to ER-Golgi contacts upon ER stress (Liu et al, 2017). Similarly, Tcb3 re-localises from contacts of the cortical ER and the plasma membrane to ER-Golgi contacts (Ikeda et al al, 2020). Indeed, tunicamycin treatment triggered co-localisation of Nvj2 and Tcb3 with Snf7-positive ER subdomains (Figure 7A, B). This phenotype was particularly clear for Tcb3. More than 90% of ER-localised Snf7 puncta contained Tcb3, and 60% additionally contained Cop1 (Figure 7C, D). To determine whether re-localisation of Tcb3 depended on ER association of ESCRT proteins, we quantified stress-induced Tcb3 puncta at cytoplasmic ER clusters in wild-type or bro1(Y320D) cells. The number of such puncta distinct from Tcb3 signal at the cell cortex was strongly reduced in bro1(Y320D) cells (Figure 7E, F). We then tested whether Tcb3 and Nvj2 were needed to establish Snf7-containing ER-Golgi contacts by analysing ER recruitment of Snf7 as well as co-localisation of Cop1 and Snf7. However, even combined removal of Nvj2, Tcb3 and its homologues Tcb1 and Tcb2 had no discernable effect (Figure S6A, B). Hence, different or additional tethers maintained stress-induced ER-Golgi contacts.

**Figure 7.**
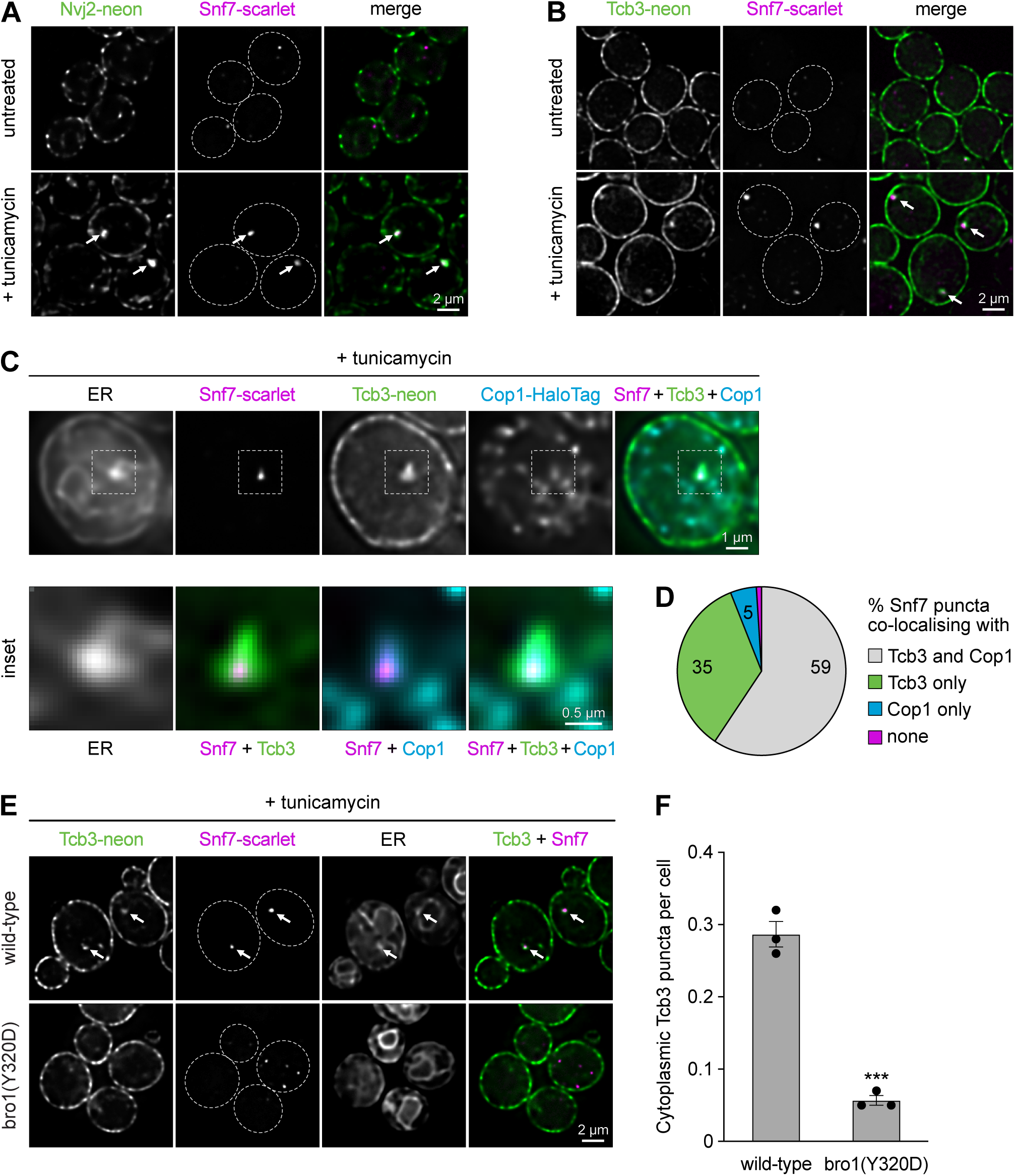
Stress-induced ER-Golgi contacts contain lipid transfer proteins. **(A)** Deconvolved fluorescent images of untreated and tunicamycin-treated wild-type cells expressing Nvj2-neon, Snf7-scarlet and the ER marker BFP-Ubc6. Dotted lines indicate cell boundaries. Arrows mark Snf7-scarlet puncta. **(B)** As in panel A but of cells expressing Tcb3-neon instead of Nvj2-neon. **(C)** Deconvolved fluorescent images of tunicamycin-treated cells expressing BFP-Ubc6, Snf7-scarlet, Tcb3-neon and the COPI protein Cop1-HaloTag. The area marked in the upper panel is shown at larger magnification in the lower panel. **(D)** Quantification of Snf7 puncta as shown in panel C that contained Snf7+Tcb3+Cop1, Snf7+Tcb3, Snf7+Cop1 or only Snf7. A total of 101 ER-localised Snf7-scarlet puncta were assessed for co-localisation with Tcb3 and Cop1. **(E)** Deconvolved fluorescent images of tunicamycin-treated wild-type and bro1(Y320D) cells expressing Tcb3-neon, Snf7-scarlet and BFP-Ubc6. Dotted lines indicate cell boundaries. Arrows mark Snf7-scarlet puncta. **(F)** Quantification of cytoplasmic Tcb3 puncta in tunicamycin-treated WT and bro1(Y320D) cells. Only Tcb3 structures separate from the signal at the cell cortex were counted. Data are the mean of n = 3 biological replicates, error bars indicate the standard error of the mean. Statistical significance of the difference between wild-type and bro1(Y320D) cells was evaluated with a two-tailed homoscedastic t-test. *** p<0.001.

These results showed that Snf7 puncta at ER subdomains coincide with Golgi structures and the ER-localised tethering proteins Nvj2 and Tcb3, representing a stress-inducible ER-Golgi contact. Tcb3 localisation to these contacts relied on ER recruitment of Bro1 and, presumably, on other ESCRT proteins such as Snf7.

### ESCRT-dependent ER-Golgi contacts contribute to cell fitness

Both Nvj2 and Tcb3 can transfer ceramide between ER and Golgi (Liu et al, 2017; Ikeda et al, 2020). Ceramide is synthesised in the ER, is transported to the Golgi by vesicular and non-vesicular routes, and is converted into complex sphingolipids by Golgi-resident enzymes (Schlarmann et al, 2021; Clausmeyer and Fröhlich, 2023). During ER stress, ceramide can accumulate in the ER and be toxic (Epstein et al, 2012; Liu et al, 2017; Yabuki et al, 2019). Besides export to the Golgi, another pathway for eliminating ceramide is its conversion into acylceramide and deposition in lipid droplets (Voynova et al, 2012). Disruption of both pathways by combined removal of Nvj2 and the two acyltransferases for acylceramide synthesis, Dga1 and Lro1, impairs cell fitness (Liu et al, 2017). To test if disruption of ESCRT recruitment to the ER had similar effects, we compared the growth of wild-type, *bro1(Y320D)*, *dga1Δ lro1Δ* and *bro1(Y320D) dga1Δ lro1Δ* cells. Only *bro1(Y320D) dga1Δ lro1Δ* cells showed a growth defect (Figure 8A). Thus, ESCRT recruitment to the ER becomes important for fitness when Dga1 and Lro1 are absent. Dga1 and Lro1 convert ceramide into acylceramide and also diacylglycerol into triacylglycerol. Hence, accumulation of ceramide and diacylglycerol may both have contributed to the observed phenotype.

**Figure 8.**
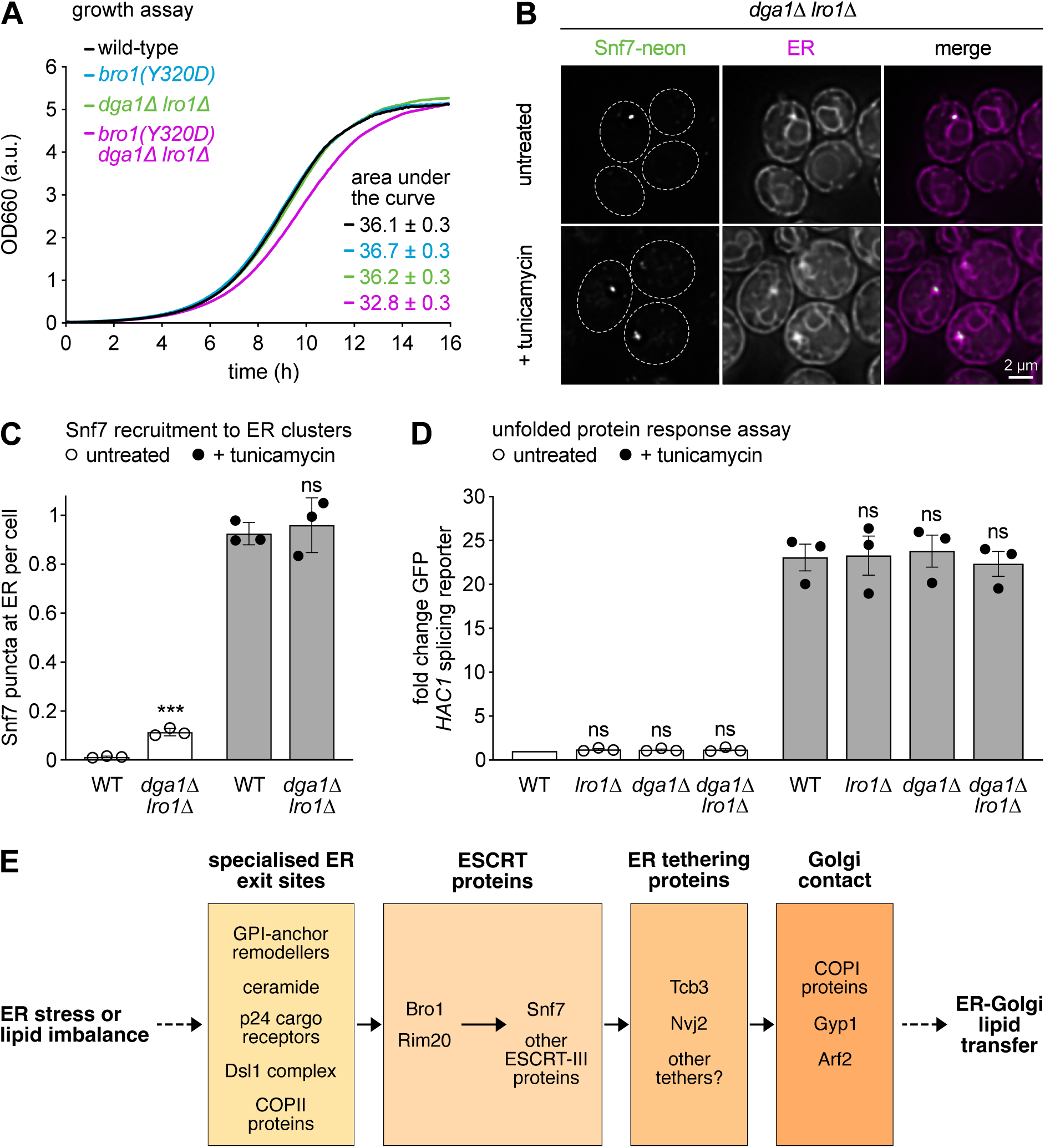
ESCRT-dependent ER-Golgi contacts contribute to cell fitness. **(A)** Growth assays of the indicated strains in standard liquid growth medium. Plotted is the optical density of growing cultures at 660 nm (OD660) in arbitrary units (a.u.). Data are the mean of n = 3 biological replicates. The area under the curve is given as mean ± standard error of the mean. **(B)** Deconvolved fluorescent images of untreated and tunicamycin-treated *dga1*Δ *lro1*Δ cells expressing Snf7-neon and the ER marker cherry-Ubc6. Dotted lines indicate cell boundaries. **(C)** Quantification of Snf7-neon puncta at ER clusters per cell in untreated and tunicamycin-treated wild-type (WT) and *dga1*Δ *lro1*Δ cells expressing Snf7-neon and cherry-Ubc6. Data are the mean of n = 3 biological replicates, error bars indicate the standard error of the mean. Statistical significance of the difference between wild-type and *dga1Δ lro1Δ* was evaluated with a two-tailed homoscedastic t-test. *** p<0.001, ns = not significant. **(D)** Fold change GFP fluorescence arising from the *HAC1* splicing reporter in untreated and tunicamycin-treated wild-type, *lro1Δ*, *dga1Δ* and *dga1*Δ *lro1*Δ cells. Data are the mean of n = 3 biological replicates, error bars indicate the standard error of the mean. Data were normalised to the untreated wild-type. Statistical significance of the difference between the wild-type and each mutant strain was evaluated with a two-tailed t-test (heteroscedastic t-test for untreated cells, homoscedastic t-test for tunicamycin-treated cells). **(E)** Model of the formation of ESCRT-dependent ER-Golgi contacts via a recruitment cascade. ER stress or lipid imbalance cause coalescence of specialised ER exit sites for the export of GPI-anchored proteins. The integrity of this ER subdomain depends on GPI-anchor remodellers, ceramide, p24 family cargo receptors, the Dsl1 tethering complex and COPII proteins. The ESCRT-associated proteins Bro1 and Rim20 localise to this ER subdomain and recruit Snf7 and other ESCRT-III proteins. ESCRT proteins are needed to recruit the tethering and lipid transfer protein Tcb3. Nvj2 is enriched at the same sites. Contact formation with the Golgi requires COPI proteins, which are enriched, together with Gyp1 and Arf2, at the Golgi subdomain contacting the ER. ER-Golgi contact still occurs in the absence of Tcb1/2/3 and Nvj2 so that additional tethers must be involved. The generation of ER-Golgi contact promotes lipid transfer between the ER and Golgi to maintain organelle homeostasis.

Remarkably, removal of Dga1 and Lro1 by itself triggered the formation of ER-localised Snf7 puncta in approximately 10% of cells (Figure 8B, C). Removal of Dga1 and Lro1 by itself caused only minimal activation of the unfolded protein response (Figure 8D). These levels of stress signalling did not induce Snf7 puncta when elicited by low concentrations of tunicamycin (Figure S6C). Hence, loss of Dga1 and Lro1 likely induced Snf7 puncta not by causing protein misfolding and activating the unfolded protein response but by imbalancing lipid metabolism. These results indicated that cells respond to perturbed lipid homeostasis by forming ESCRT-dependent ER-Golgi contacts, which contribute to cell fitness.

## Discussion

We report that ER stress triggers recruitment of ESCRT-III proteins to stress-induced ER-Golgi contacts. Contact formation involves specialised ER exit sites known to consist of ceramide-rich membrane domains. ESCRT-III proteins are recruited to these ER subdomains by two Bro1 domain proteins and help gather ER tethering and lipid transfer proteins (Figure 8E). Loss of ESCRT recruitment impairs growth of cells with perturbed lipid metabolism, showing that ESCRT-dependent ER-Golgi contacts help maintain cell homeostasis. Our findings indicate that specialised ER exit sites can be repurposed for contacting the Golgi and reveal a function of ESCRT proteins in organising a stress-inducible organelle contact. These advances raise many new questions about the mechanisms of ER subdomain formation, ESCRT recruitment to the ER, the architecture of ESCRT-dependent ER-Golgi contacts, and their functions.

First, how does ER stress induce ER subdomains derived from specialised ER exit sites? Many newly-synthesised secretory proteins misfold during stress and are retained in the ER, especially GPI-anchored proteins (Platzek et al, 2025). Their accumulation in the ER may induce rearrangement of the ER exit sites that normally mediate their export, as reflected by enrichment of the cargo receptor Emp24 and COPII proteins at ER clusters. Accordingly, ER clustering was diminished upon disruption of GPI anchor remodelling, sphingolipid synthesis, cargo capture by p24 family receptors or COPII function. Curiously, cells mostly had only a single Snf7-containing ER cluster. General inhibition of ER export causes coalescence of ER exit sites (Shindiapina and Barlowe, 2010; Okamoto et al, 2012; Schröter et al, 2016), and a similar scenario may play out for specialised ER exit sites during ER stress.

Second, how do ESCRTs associate with the ER? Recruitment of the ESCRT-III protein Snf7 was independent of ESCRT-0/I/II but required its interactors Bro1 and Rim20. The human Bro1 homologue ALIX can associate with membranes by binding to lysobisphosphatidic acid, but the lipid binding motif in the Bro1 domain of ALIX is not present in yeast Bro1 (Bissig et al, 2013). ALIX also interacts with ALG2, which can bind membranes and collaborates with ALIX and ESCRT-III in microautophagy of ER exit sites (Shukla et al, 2024; Liao et al, 2024). However, ALIX-ALG2 interaction and microautophagy of ER exit sites require the proline-rich domain of ALIX (Shibata et al, 2004; Liao et al, 2024). Bro1 recruitment to the ER did not depend on its proline-rich domain. Moreover, Bro1 recruitment did not involve the ALG2 homologue Pef1 (unpublished results). Intriguingly, ER targeting of Bro1 did require a hydrophobic surface patch in its Bro1 domain, which is centred around tyrosine-320, is conserved in Rim20 and ALIX, and likely functions as interaction interface (Kim et al, 2005). The identity of its putative binding partner remains to be discovered. Furthermore, tunicamycin induced ER recruitment of Snf7 much more efficiently than dithiothreitol, even though both drugs cause strong ER stress (Platzek et al, 2025). Hence, tunicamycin-specific effects, perhaps involving lipid alterations, may play a role.

Third, what is the architecture of ESCRT-dependent ER-Golgi contacts? The ER membrane at these sites was convoluted and likely bound by several ESCRT proteins. These ESCRTs may remodel the ER membrane and contribute to its peculiar morphology. ESCRTs usually associate with membranes only transiently but they can also form stable scaffolds (Penalva, 2014; Stempels et al, 2023). We expressed tagged Snf7 at low levels alongside endogenous Snf7 and found that it did not disturb endosomal sorting and required endogenous Snf7 for ER localization. However, it was not a neutral tracer. Tagged Snf7 stabilised Bro1 and Rim20 at ER-Golgi contacts, which aided microscopic analysis but likely obscured ESCRT dynamics. Better tools are needed to visualise ESCRT assembly and disassembly at ER-Golgi contacts and resolve whether ESCRTs act as scaffolds. In mammals, COPII proteins have been proposed to form collars at the base of tubular ER extension that protrude towards the Golgi, and COPI and COPII proteins bind to distinct regions of these extensions (Weigel et al, 2021; Shomron et al, 2021). Our data show that COPI and II proteins also have distinct distributions in proximity of ER-localised ESCRT assemblies. Nvj2 and Tcb3 at stress-induced ER-Golgi contacts have been reported to co-localise with the medial-Golgi (Liu et al, 2017; Ikeda et al, 2020). We found extensive co-localisation of Snf7 with COPI proteins, Arf2 and especially Gyp1, but only moderate overlap with other Golgi markers. The yeast Golgi is not stacked and individual cisternae undergo maturation (Losev et al, 2006; Matsuura-Tokita et al, 2006). Gyp1 marks Golgi cisternae throughout their maturation because its Golgi association begins at an early stage and continues until a late stage (Thomas et al, 2021). The strong co-localisation of Snf7 with Gyp1 and the moderate co-localisation with markers of different maturation stages likely indicates that ER-Golgi contacts form with the early Golgi and remain in place as the Golgi matures. Alternatively, Gyp1 could be an integral part of ER-Golgi contacts. The identity of the tethers critical for these contacts remains unknown. Contacts persisted in cells lacking Nvj2 and Tcb1/2/3, implying the involvement of other or additional relevant tethers. Redundancy between tethers can be extensive. For example, ER-plasma membrane contacts in yeast are mediated by at least six proteins (Manford et al, 2012).

Fourth, what are the functions of ESCRT-dependent ER-Golgi contacts? ESCRT recruitment enabled enrichment of Tcb3 at these sites, potentially promoting non-vesicular lipid transport. Tcb3 is related to human extended synaptotagmin proteins and prefers high-curvature ER membrane, as is present at ER-Golgi contacts (Manford et al, 2012; Hoffmann et al, 2019). Nvj2 and Tcb3 transfer ceramide from the ER to the Golgi during ER stress (Liu et al, 2017; Ikeda et al, 2020). Since stress-induced ER-Golgi contacts derive from specialised ER exit sites consisting of ceramide-rich membrane, the local ceramide concentration should be high. Furthermore, ceramide levels in the ER rise during stress, possibly due to reduced vesicular transport to the Golgi, and they pose a threat to proper ER function (Epstein et al, 2012; Liu et al, 2017; Yabuki et al, 2019; Hwang et al, 2023). Non-vesicular transport via inducible ER-Golgi contacts therefore is a mechanism to avert lipotoxicity. A second mechanism for alleviating lipotoxicity is the synthesis of acylceramide and triacylglyceride, and their deposition in lipid droplets (Voynova et al, 2012). Accordingly, simultaneous loss of ESCRT recruitment to the ER and the acyltransferases for acylceramide and triacylglyceride synthesis caused a growth defect. Interestingly, this growth defect was observed even in otherwise unstressed cells, and blocking acylceramide and triacylglyceride synthesis by itself was sufficient to trigger ESCRT recruitment to ER-Golgi contacts. Thus, ESCRT recruitment may not be limited to ER stress conditions. This notion is supported by the observation that Snf7 recruitment can also be induced by Bro1 or Rim20 overexpression. It remains to be explored whether ESCRT recruitment may even occur constitutively.

The utilisation of specialised ER exit sites for the formation of ER-Golgi contacts is a previously unknown aspect of the cellular response to ER stress. ESCRTs participate in many membrane remodelling processes, and their role in organising organelle contacts extends the remarkable scope of the ESCRT machinery even further.

## Materials and Methods

### Plasmids

Plasmids used in this study are listed in Table S4. Plasmids of the pFA6a series have been described (Longtine et al, 1998, Janke et al, 2004; Young et al, 2013; Larochelle et al, 2019; Papagiannidis et al, 2021; Platzek et al, 2025; Hubbe et al, 2026). To generate pRS416-P_VPS24_-Snf7-LAP-mScarlet-i, mScarlet-i was amplified from pFA6a-mScarlet-i-kanMX6 and combined by Gibson assembly with pRS416-P_VPS24_-Snf7-LAP-mNeonGreen (Schäfer et al, 2020), replacing mNeonGreen. To generate pRS413/5-P_VPS24_-Snf7-LAP-mScarlet-i, P_VPS24_-Snf7-LAP-mScarlet-i was excised from pRS416-P_VPS24_-Snf7-LAP-mScarlet-i with NaeI/SacI and inserted into the NaeI/SacI sites of pRS413-P_CYC_ or pRS415-P_CYC_ (Mumberg et al, 1995). To generate pRS304-P_TEF_-TagBFP-Ubc6, P_TEF_-TagBFP-Ubc6 was excised from pRS406-P_TEF_-TagBFP-Ubc6 (Platzek et al, 2025) with SacI/KpnI and inserted into the SacI/KpnI site of pRS304 (Sikorski and Hieter, 1989). To generate pRS404-P_GPD_-TagBFP-Ubc6, the *GPD* promoter was excised from pRS404-P_GPD_-mCherry-Ubc6 (Schäfer et al, 2020) with SacI/SpeI and inserted into the SacI/SpeI site of pRS404-P_TEF_-TagBFP-Ubc6, replacing the *TEF* promoter. To generate pRS315-Ire1-HA, Ire1-HA including upstream and downstream sequence was excised from pRS316-Ire1-HA with NotI/XhoI and cloned into the NotI/XhoI site of pRS315 (Sikorski and Hieter, 1989). To generate pRS315-Ire1(mScarlet-i)-HA, pRS315-Ire1-HA was linearised by PCR and combined by Gibson assembly with mScarlet-i amplified from pFA6a-mScarlet-i-natMX6 such that mScarlet-i was inserted between residues I571 and G572 of Ire1. To generate pRS413-Bro1, the *BRO1* gene including promoter and terminator was amplified from genomic DNA and combined with pRS413-P_ADH_ (Mumberg et al, 1995) by Gibson assembly, replacing the *ADH* promoter and *CYC* terminator. Site-directed mutagenesis was then used to introduce point mutations into pRS413-Bro1 and thus generate pRS413-bro1(K246A) and pRS413-bro1(Y320D).

### Yeast strain generation and culture

Unless indicated otherwise, strains in this study were derived from strain SSY122 in the W303 background (genotype *ADE2 leu2-3,112 trp1-1 ura3-1 his3-11,15 MATa*) and are listed in Table S5. Gene tagging, gene deletion and promoter exchange was done with PCR products (Longtine et al, 1998; Janke et al., 2004). Strain SSY4470 expressing C-terminally truncated bro1(1-387) was generated by a partial deletion of the *BRO1* open reading frame. Point mutations were introduced into the endogenous *BRO1* gene locus by a two-step procedure. First, the *URA3* gene was amplified from plasmid pFA6a-URA3 and integrated between nucleotide 434 and 963 of the *BRO1* coding sequence. Second, bro1(K246A) or bro1(Y320D) were amplified from plasmid pRS413-bro1(K246A) or pRS413-bro1(Y320D) and integrated into the disrupted *BRO1* coding sequence such that they replaced the *URA3* gene, restored the coding sequence and introduced the respective point mutation. Clones that had lost the *URA3* gene were selected on plates containing 5’-fluoroorotic acid, and the point mutations were confirmed by sequencing. To introduce the F196A, F197A double mutation into the C-terminal region of Emp24, the *EMP24* gene was tagged with mNeonGreen as above except that the forward primer used to generate the necessary PCR product contained point mutations, which, upon integration of the PCR product into the *EMP24* locus, converted the codons for F196 and F197 into codons for alanines. For genomic integration of centromeric pRS41X plasmids, the expression cassettes including the selection markers were amplified by PCR such that ends homologous to the genomic target loci were added to the PCR product. For genomic integration of the integrative plasmids pSS349 and pDK45, the expression cassettes including the selection markers were excised with AfeI/NaeI and PsiI, respectively, before transformation. Unless indicated otherwise, strains were grown in liquid culture at 30°C in synthetic complete medium containing 0.7% yeast nitrogen base (Merck), amino acids and 2% glucose (SCD medium).

### Western blotting

Cultures were grown to mid log phase (OD_600_ = 0.5) in SCD medium, cells were collected by centrifugation, washed once with water, resuspended in lysis buffer (50 mM HEPES pH 7.5, 0.5 mM EDTA, Roche complete protease inhibitors), and disrupted by bead beating with a FastPrep-24 (MP Biomedicals). Proteins were solubilised by addition of 1.5% w/v SDS and incubation at 65°C for 5 min. Lysates were cleared at 16,000 × g at 4°C for 2 min, and protein concentrations were determined with a Pierce BCA protein assay kit (Thermo Fisher Scientific). All subsequent steps were done as described (Hubbe et al, 2026). Primary antibodies were rabbit anti-Snf7 (Heinzle et al, 2019), mouse anti-myc (clone 9B11 from Cell Signaling Technology, AB_331783), mouse anti-FLAG (clone M2 from Sigma, AB_262044) and mouse anti-Pgk1 (clone 22C5D8 from Abcam, AB_10861977). Secondary antibodies were goat anti-rabbit Alexa-680 (Invitrogen, AB_2535736), goat anti-mouse Alexa-680 (Invitrogen, AB_141436) and goat anti-mouse IRDye-800CW (LI-COR, AB_621842). Fluorescence was detected with an Odyssey CLx imaging system (LI-COR).

### Endosomal sorting assay

Strains expressing Mup1-pHluorin (Ivashov et al, 2020) along with cytosolic BFP and a control strain only expressing cytosolic BFP were grown in 1 ml SCD medium without methionine in 96-deep well plates for 24 h so that cultures reached saturation. These pre-cultures were diluted into 1 ml fresh methionine-free SCD medium and grown overnight so that they reached mid log phase (OD_600_ = 0.1 – 0.5). Overnight cultures were diluted to OD_600_ = 0.05 in methionine-free SCD medium in duplicate 1-ml cultures. One culture per strain was left untreated, the other culture was supplemented with 20 µg/ml methionine to induce endocytosis of Mup1-pHluorin. Total cell fluorescence after excitation with a 405 nm laser (blue fluorescence) or a 488 nm laser (green fluorescence) was measured with a FACSCanto flow cytometer (BD Biosciences) at 10 minute intervals for up to 90 minutes. The geometric mean of blue and green fluorescence per cell was calculated with FlowJo software. Cell autofluorescence after excitation with the 488 nm laser was determined with the control strain and subtracted from the green fluorescence of the remaining strains. Cellular Mup1-pHluorin levels outside the vacuole were then determined by dividing the background-corrected green fluorescence by the blue fluorescence, which served as a measure for cell size. Finally, this pHluorin/BFP ratio in methionine-treated cells was divided by the ratio in untreated cells to determine methionine-induced quenching of Mup1-pHluorin fluorescence. Data were normalised to the Mup1-pHluorin fluorescence at t = 0.

### Microscopy

Unless indicated otherwise, cultures grown to OD_600_ = 0.3 in SCD medium were treated with 1 µg/ml tunicamycin (Calbiochem) or 8 mM DTT (Roche) for 3 h. Cells from 1 ml culture were pelleted and resuspended in 20 µl medium. Three microliters cell suspension were spotted onto cover slips, covered with an agarose pad prepared with SCD medium and imaged with an Olympus IX81 CellSens microscope equipped with a PLAPO 100x/1.45 objective and a Hamamatsu Orca R2 camera (images in Figure S1C) or a Nikon Ti2 microscope equipped with a PLAN APO 100x/1.45 oil objective and a Hamamatsu Orca Fusion-BT camera (all other images). For image quantification, Z-stacks of ten optical slices with a spacing of 1 µm were acquired to capture the entire volumes of all cells in a field of view. For subsequent image deconvolution, Z-stacks of eleven optical slices with a spacing of 0.2 µm were acquired and processed with the Richardson-Lucy algorithm (30 iterations, automatic detection of noise level) using the NIS Elements Software deconvolution module (Nikon). Deconvolution reduced out of focus light at the expense of comparability of signal intensity across images. Even though images were acquired in a way that avoided pixel saturation, deconvolved images contained saturated pixels and can therefore only be compared qualitatively.

### Image quantification

For manual quantification of ER clusters, Snf7 puncta at ER clusters, Bro1 puncta at ER clusters and Tcb3 puncta (Figures 1B, 2A, 2B, 2D, 3C, 3D, 5A, 5B, 5D, 5F, 7D, 7F, 8C, S2C, S2E, S6C), Z-stacks were converted to Z-max intensity projections and anonymised with the “Blind Analysis Tools” plug-in in ImageJ (https://imagej.net/plugins/blind-analysis-tools) before analysis to prevent user bias. For automated quantification of Snf7-neon puncta at ER clusters (all other quantifications), cells were analysed using a custom Python script (https://github.com/SchuckLab/Snf7-puncta-counter). The script first uses intensity thresholding of the brightfield image to mask cell areas. It then uses adaptive thresholding to detect bright Snf7-neon structures in maximum intensity projections of Z-stacks and assesses the circularity and size of each structure to exclude bright patches that are dissimilar in shape to Snf7-neon puncta. Each structure is then assigned to the corresponding cell mask to obtain the percentage of cells with Snf7-neon puncta and the average number of Snf7-neon puncta per cell. In both, manual and automated quantification, at least 150 cells were analysed per condition and biological replicate.

Co-localisation of Snf7 with COPI or Golgi proteins was scored by sample anonymisation as above and visual assessment of signal overlap at 50 ER-localised Snf7 puncta. Margins of error at the 95% confidence level in Figure 6C, D and F were calculated by the modified Wald method using the GraphPad website (https://www.graphpad.com/quickcalcs/confinterval1).

### Light microscopy-guided cryo-electron tomography

To collect tomograms of Snf7-containing ER clusters, strain SSY3786 was used, which expressed Snf7-scarlet, BFP-Ubc6 and Sec24-neon, and lacked Chm7. Removal of Chm7 had no significant impact on the formation of ER-localised Snf7 puncta (Figure 2A) but prevented Snf7 accumulation at sites of nuclear pore complex quality control, which proved distracting during subsequent light microscopy-guided focussed ion beam (FIB) milling of the sample. Cells were grown to OD_600_ = 0.7 in SCD medium and treated with 1 µg/ml tunicamycin for 3 h. Dynabeads MyOne Silane (Thermo Fisher Scientific) were added to yeast culture at a 1:15 dilution, and 3.5 µl culture was incubated on R1/2 200 mesh glow discharged copper grids (Quantifoil) for 30 seconds and plunge-frozen in liquid ethane/propane (37%/63%). Lamellae were prepared using an Aquilos cryo-FIB/SEM (Thermo Fisher Scientific) with an integrated METEOR fluorescence microscope (Delmic). Micro-expansion joints were added before milling to increase lamella stability. Automated milling was carried out using the AutoLamella Python package (Wagner et al, 2020) and done in steps of decreasing current and thickness (Table S6). After the third step, fluorescence images were acquired with an Olympus LMPLFLN 50x/0.8 air objective, a Zyla Andor 4.2 sCMOS camera and Odemis 3.2.1 software. Lamellae still showing Snf7-scarlet fluorescence were manually polished with a current of 30 pA for up to 7 minutes to reach a final thickness of approximately 200 nm.

As preparation for light microscopy-guided cryo-ET data collection, SEM images of the polished lamellae were overlayed with the corresponding fluorescence image. A standard scaling factor (0.583 ± 0.003, n = 11) was obtained with the 3D correlation toolbox (Arnold et al, 2016) and applied to the fluorescence image in FIJI software to match it to the 1,000x SEM images. The scaled fluorescence image was placed on top of the SEM image, and rotated and translated to create the final overlay using Dynabeads and grid holes as reference points. Lamellae were then imaged on a 300 kV FEI Krios TEM (Thermo Fisher Scientific) with a K3 summit direct electron detector (Gatan), equipped with a post-column energy filter aligned to the zero-loss peak and a 20 keV slit width. Overview images were taken with SerialEM (Mastronarde, 2003) at 6,500x magnification (28.6 Å/pixel). These TEM overviews were manually aligned to the overlay of fluorescence and SEM images. Tilt series were collected at locations with clearly visible Snf7 puncta. Tilt series were recorded at a pixel size of 3.29 Å/pixel, a dose rate of about 15 e/pixel/s, and a total dose of approximately 100 e/Å^2^. Tilt series were collected with a tilt increment of 3°, a defocus target of -4 µm and a tilt range of approximately 66° to -54° or 54° to -66°, depending on lamella orientation.

Tilt series movies were motion corrected using MotionCor2 (Zheng et al, 2017), low-quality tilts were excluded, and aligned stacks were generated in WARP (Tegunov and Cramer, 2019). Pre-processed tilt stacks were aligned to a common 3D coordinate system and reconstructed in AreTomo (Zheng et al, 2022), tomograms were denoised with CryoCARE v0.3.0 (Buchholz et al, 2019) and membranes were detected using MemBrainSeg (6x downsampled, 19.74 Å/pixel; Lamm et al, 2024). Segmentations were inspected and manually polished in Avizo v9.2.0 (FEI). To visualise ER-bound ribosomes, template matching was performed with pytom-match-pick (Chaillet et al, 2023) on 6x downsampled tomograms using an 80S yeast ribosome (EMDB-18231) as template and a mask with a radius of nine pixels (corresponding to 177.66 Å). To remove cytosolic ribosomes, a mask surrounding the ER was created by dilating the ER segmentations 15 times in Avizo and applied to the obtained particle coordinates. Template matching results were evaluated using the ArtiaX plugin in ChimeraX, tomograms were visualised in IMOD, and segmentations and ribosomes were visualised in ArtiaX (Mastronarde and Held, 2017; Ermel et al, 2022).

### Snf7 biotinylation assay

Cells expressing Snf7-neon and TurboID-tagged Bro1 variants in a *vps27Δ* background were grown to early log phase in SCD medium and left untreated or treated with 1 µg/ml tunicamycin for 3 h. No biotin was added besides the 2 µg/l biotin already present in SCD medium. Fifty ODs of cells were harvested by centrifugation, resuspended in lysis buffer (50 mM HEPES pH 7.5, 0.5 mM EDTA, Roche complete protease inhibitors, 1 mM PMSF, 0.3% w/v SDS) and lysed by bead beating. For each sample, 15 µl high capacity streptavidin agarose beads (Thermo Fisher Scientific) washed twice with 200 µl lysis buffer were used, and biotinylated proteins were precipitated from 1 mg total cell protein in 500 µl lysis buffer at 4°C for 3 h. Beads were washed at 4°C for 2 x 5 min with 500 µl wash buffer 1 (50 mM HEPES pH 7.5, 0.5 mM EDTA, Roche complete protease inhibitors, 1 mM PMSF) and for 2 x 5 min with 500 µl wash buffer 2 (50 mM HEPES pH 7.5, 0.5 mM EDTA). Bound biotinylated protein was eluted with 25 µl SDS-PAGE sample buffer containing 0.1 volumes ß-mercaptoethanol and 2 mM biotin at 65°C for 10 min and analysed by western blotting.

### Biotinylation screen

Cells expressing Snf7-neon and TurboID-tagged Bro1 or bro1(K246A) in a *vps27Δ chm7Δ pep4Δ prb1Δ* background were grown to early log phase in SCD medium and left untreated or treated with 1 µg/ml tunicamycin for 3 h. No biotin was added besides the 2 µg/l biotin already present in SCD medium. Five hundred ODs of cells were harvested by centrifugation, resuspended in lysis buffer (50 mM Tris-HCl pH 7.5, 150 mM NaCl, 5 mM EDTA, Roche complete protease inhibitors, 1 mM DTT, 0.4% w/v SDS, 2% w/v TX-100) and lysed by bead beating. For each sample, 60 µl high capacity streptavidin agarose beads (Thermo Fisher Scientific) washed twice with 1 ml lysis buffer were used, and biotinylated proteins were precipitated from 25 mg total cell protein in 5 ml lysis buffer at 4°C for 3 h. Beads were washed at room temperature for 2 x 5 min with 500 µl wash buffer 1 (50 mM Tris-HCl pH 7.5, 150 mM NaCl, 5 mM EDTA, Roche complete protease inhibitors, 1 mM DTT, 0.4% w/v SDS, 2% w/v TX-100), for 2 x 5 min with 500 µl wash buffer 2 (2% w/v SDS), for 2 x 5 min with 500 µl wash buffer 3 (50 mM HEPES pH 7.4, 500 mM NaCl, 1 mM EDTA, 1% w/v Triton X- 100, 0.1% w/v sodium deoxycholate) and for 2 x 5 min with wash buffer 4 (50 mM Tris-HCl pH 7.5, 50 mM NaCl, 0.1% w/v Triton X-100). Bound biotinylated protein was eluted with 30 µl SDS-PAGE sample buffer containing 0.1 volumes ß-mercaptoethanol and 2 mM biotin for 10 min at 65°C. The screen was done in three biological replicates.

### Mass spectrometry

Biotinylated proteins were separated by SDS-PAGE and stained with Coomassie. Lanes were excised and cut just above the band of the abundant endogenously biotinylated protein Arc1 at about 50 kDa to reduce distortion of the subsequent mass spectrometric measurements. The resulting two gel pieces per sample were processed individually. Proteins were digested in-gel with trypsin (Pedre et al, 2023), peptides were concentrated by vacuum centrifugation and dissolved in 0.1% TFA. Nanoflow LC-MS/MS analysis was performed with an Ultimate 3000 liquid chromatography system coupled to an Orbitrap QE HF (Thermo Fisher Scientific). An in-house packed analytical column (75 µm x 200 mm, 1.9 µm ReprosilPur-AQ 120 C18 material) was used to separate peptides. Mobile phase solutions were prepared with solvent A (0.1% formic acid, 1% acetonitrile) and solvent B (0.1% formic acid, 89.9% acetonitrile). Peptides from the upper part of the gel were separated with a linear gradient starting from 3% solvent B, increasing to 23% over 50 min and to 38% over 10 min, followed by washout with 95% solvent B. Peptides from the lower part of the gel were separated the same way, except that step times were 21 and 4 min. The mass spectrometer was operated in data-dependent acquisition mode. MS1 spectra (m/z 400–1600) were acquired at 60,000 (m/z 400) resolution. MS2 spectra were generated for up to 15 precursors with normalised collision energy of 27 and isolation width of 1.4 m/z.

Data analysis was done with MaxQuant 2.0.1.0 software (Cox and Mann, 2008) using the Uniprot *S. cerevisiae* protein database of reference strain ATCC 204508 from March 2022. Statistical analyses of LFQ intensities were done with Perseus software (Tyanova et al, 2016). Pre-filtering removed common contaminants, proteins only identified by site and hits to the reverse decoy database. Samples were grouped according to strain and condition, yielding four groups with three replicates each. For each pair-wise comparison, i.e. treated versus untreated wild-type Bro1 and treated versus untreated bro1(K246A), only proteins with three values in at least one condition were considered. Missing values were imputed from a normal distribution with a width of 0.3 and a 1.8 standard deviation down shift. Two-sided t-tests were carried out for tunicamycin-treated versus untreated wild-type cells and tunicamycin-treated versus untreated bro1(K246A) cells. Volcano plots were generated with a permutation-based FDR (q-value) < 0.05 (500 permutations, S0 parameter set to 0.2). Hits were defined by a positive log_2_ fold change and a q-value <0.05. 185 and 39 proteins were enriched in the proximity of Bro1 during ER stress in wild-type and bro1(K246A) cells, respectively. Subtracting shared hits from the hits in wild-type cells and discarding transposon proteins yielded 155 final hits, which were candidate proximity partners of Bro1 at the ER during stress.

### Genetic screens

Starting from strain Y8205 in the BY4741 background (Tong and Boone, 2006), strain SSY2643 was generated, which contained Snf7-neon, the general ER marker cherry-Ubc6, the plasma membrane marker BFP-PLCdelta-PH2 and deletion of the *VPS27* gene to release Snf7 from endosomes. Synthetic genetic array technology (Tong and Boone, 2006) was then used to introduce Snf7-neon, cherry-Ubc6, BFP-PLCdelta-PH2 and the *VPS27* deletion into a collection of mutants lacking genes with ER-related functions (Table S2) and a collection of mutants with temperature-sensitive alleles (Li et al, 2011).

For screening of the ER-related gene deletion mutants, strains were grown in SCD medium in 96-deep well plates at 30°C until cultures reached saturation. Cultures were diluted to OD_600_ = 0.2 and 100 µl each were transferred into glass bottom 96-well plates (Brooks Life Sciences) coated with concanavalin A. After 2 h, the medium was changed to 400 µl SC with 5 µg/ml tunicamycin. After another 3 h, cells were imaged with a Nikon Ti-E widefield microscope with a motorised stage, the Nikon perfect focus system, a 60×/1.49 oil objective, and a Flash4 Hamamatsu sCMOS camera.

For screening of the temperature-sensitive mutants, strains were grown and treated as above except that the temperature was kept at 23°C during growth and raised to 37°C upon tunicamycin addition. After 3 h of treatment, the medium was removed and cells were fixed for 10 min by addition of 200 µl phosphate-buffered saline (PBS) with 4% w/v paraformaldehyde. Cells were then washed three times with PBS and imaged as above. Strain SSY2644 lacking *BRO1* was used as a positive control in both screens. Images were processed and viewed using custom MATLAB script (Papagiannidis et al, 2021).

### Yeast growth assays

Pre-cultures were grown in SCD medium during the day and diluted in the evening to obtain logarithmically growing cells the next morning. Overnight cultures were diluted to OD_600_ = 0.05 in fresh SCD medium and duplicate 500-µl cultures were transferred into 48-well plates. Cultures were then grown in a Tecan Spark plate reader at 30°C for 16 h and the absorbance at 660 nm was measured every five minutes. In between measurements, cultures were shaken continuously. Data were plotted with GraphPad Prism software.

### Unfolded protein response assay

Activation of the unfolded protein response was measured with the *HAC1* splicing reporter, which translates Ire1 activity into the production of GFP (Pincus et al, 2010). Cells harbouring the reporter were grown to mid log phase in 1 ml medium in 96 deep-well plates, diluted to OD_600_ = 0.05 and either left untreated or treated with tunicamycin for three hours. Forward scatter (FSC) and GFP fluorescence after excitation with a 488 nm laser were measured with a FACS Canto flow cytometer (BD Biosciences, Franklin Lakes, New Jersey) equipped with a high-throughput sampler. In parallel, autofluorescence was determined with an identically grown control strain not harbouring the *HAC1* splicing reporter. Cellular GFP fluorescence was corrected for autofluorescence and normalised to cell size by division by the FSC. Data were expressed relative to the GFP/FSC ratio in untreated wild-type cells.

## Data availability

The proteomic dataset generated during the current study are available from the ProteomeXchange Consortium via the PRIDE partner repository with the dataset identifier PXD078072.

## Acknowledgements

We thank Michael Knop, Peter Walter, David Teis and Ralf Kölling for reagents, the Flow Cytometry & FACS Core Facility at the Center for Molecular Biology at Heidelberg University, Liz Miller and Imogen Binnian for assistance, and all Schookees for comments on the manuscript. This work was supported by grants SCHU 2364/3-1 (project 455429207) and SFB-1638/1 - P02 (as part of project 511488495) from the German Research Foundation (DFG) to SS. LA additionally acknowledges support from the Luxembourg National Research Fund (project 15683139). FFr was supported by the Heisenberg programme of the DFG (project 491484150). The authors gratefully acknowledge the data storage service SDS@hd supported by the Ministry of Science, Research and the Arts Baden-Württemberg (MWK) and the DFG through grant INST 35/1503-1 FUGG.

## Author contributions

Conceptualization, L.A., O.P., J.A.S., S.S.; Software, K.O.; Formal Analysis, G.H.H.B., M.L.; Supervision, F.Fö., F.Fr., S.S.; Investigation, L.A., B.E., N.F., C.M.d.H., L.d.J., D.P., O.P., J.A.S.; Writing – Original Draft, L.A., O.P., S.S.; Writing – Review & Editing, all authors.

---

**Figure S1.**
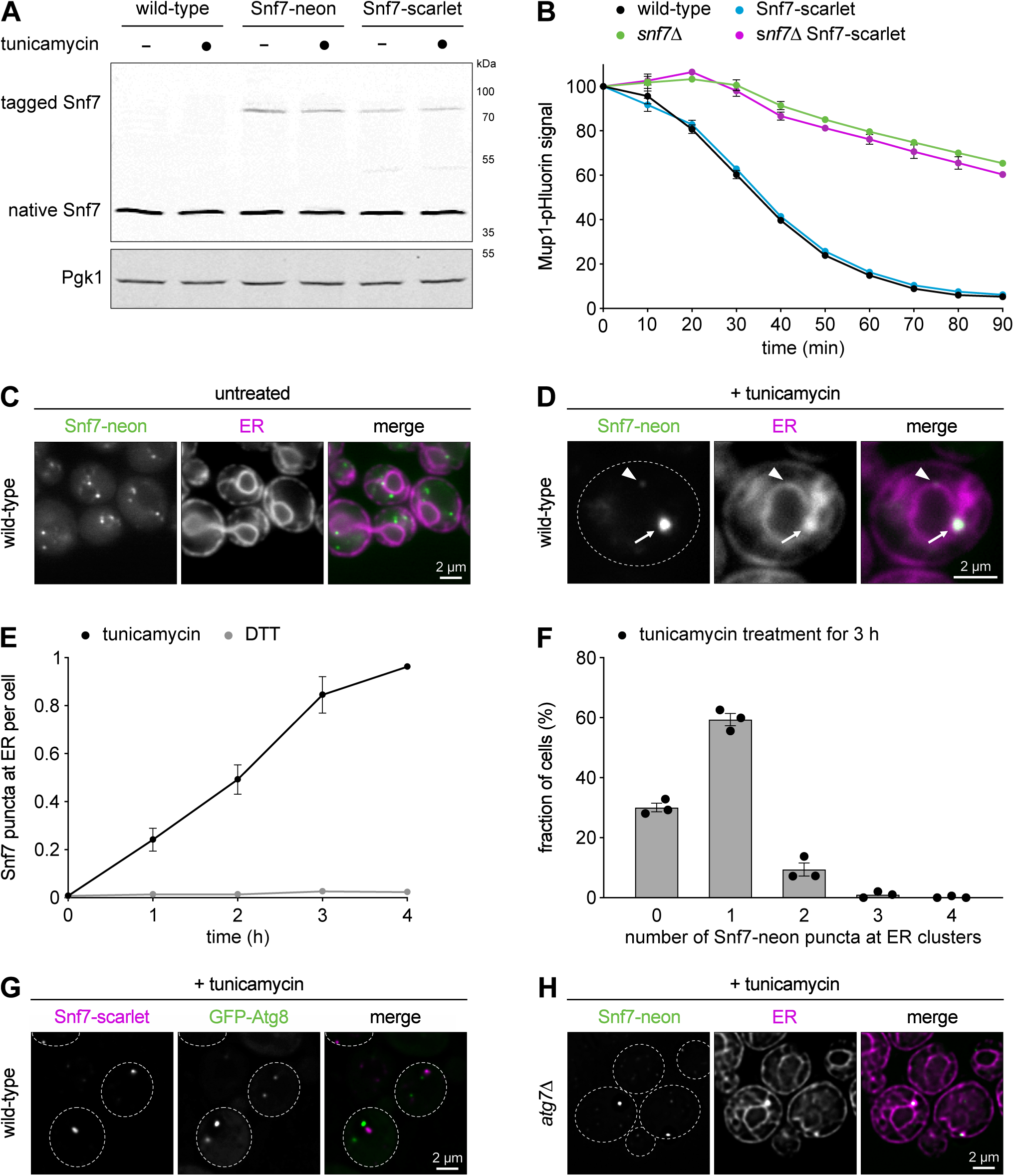
The ESCRT-III protein Snf7 is recruited to stress-induced ER subdomains. **(A)** Western blot of Snf7 from wild-type cells not expressing tagged Snf7 and from cells expressing Snf7-neon or Snf7-scarlet under the control of the *VPS24* promoter. Pgk1 served as a loading control. **(B)** Endosomal sorting assay based on Mup1-pHluorin fluorescence, as measured by flow cytometry. The x-axis indicates the time after addition of methionine to induce ESCRT-dependent transport of Mup1-pHluorin to the vacuole, where the pHluorin signal is quenched. For each strain, fluorescence was normalised to t = 0. Data are the mean of n = 3 biological replicates, error bars indicate the standard error of the mean. **(C)** Widefield fluorescent images of untreated cells expressing Snf7-neon and the ER marker cherry-Ubc6. Snf7 localises to the cytosol and to endosomes. **(D)** Widefield fluorescent images of tunicamycin-treated cells expressing Snf7-neon and the ER marker cherry-Ubc6. The dotted line indicates the cell boundary. The arrow marks a bright Snf7-neon punctum at an ER cluster, the arrowhead marks a dim Snf7-neon punctum at the nuclear envelope. **(E)** Quantification of Snf7-neon puncta at ER clusters per cell in cells treated with tunicamycin or DTT for the times indicated. Data are the mean of n = 3 biological replicates, error bars indicate the standard error of the mean. **(F)** Quantification of cells with zero to four ER-localised Snf7-neon puncta after tunicamycin treatment for 3 h. Data are the mean of n = 3 biological replicates, error bars indicate the standard error of the mean. **(G)** Deconvolved fluorescent images of tunicamycin-treated wild-type cells expressing Snf7-neon and GFP-Atg8. Dotted lines indicate cell boundaries. **(H)** Deconvolved fluorescent images of tunicamycin-treated *atg7Δ* cells expressing Snf7-neon and the ER marker cherry-Ubc6. Dotted lines indicate cell boundaries.

**Figure S2.**
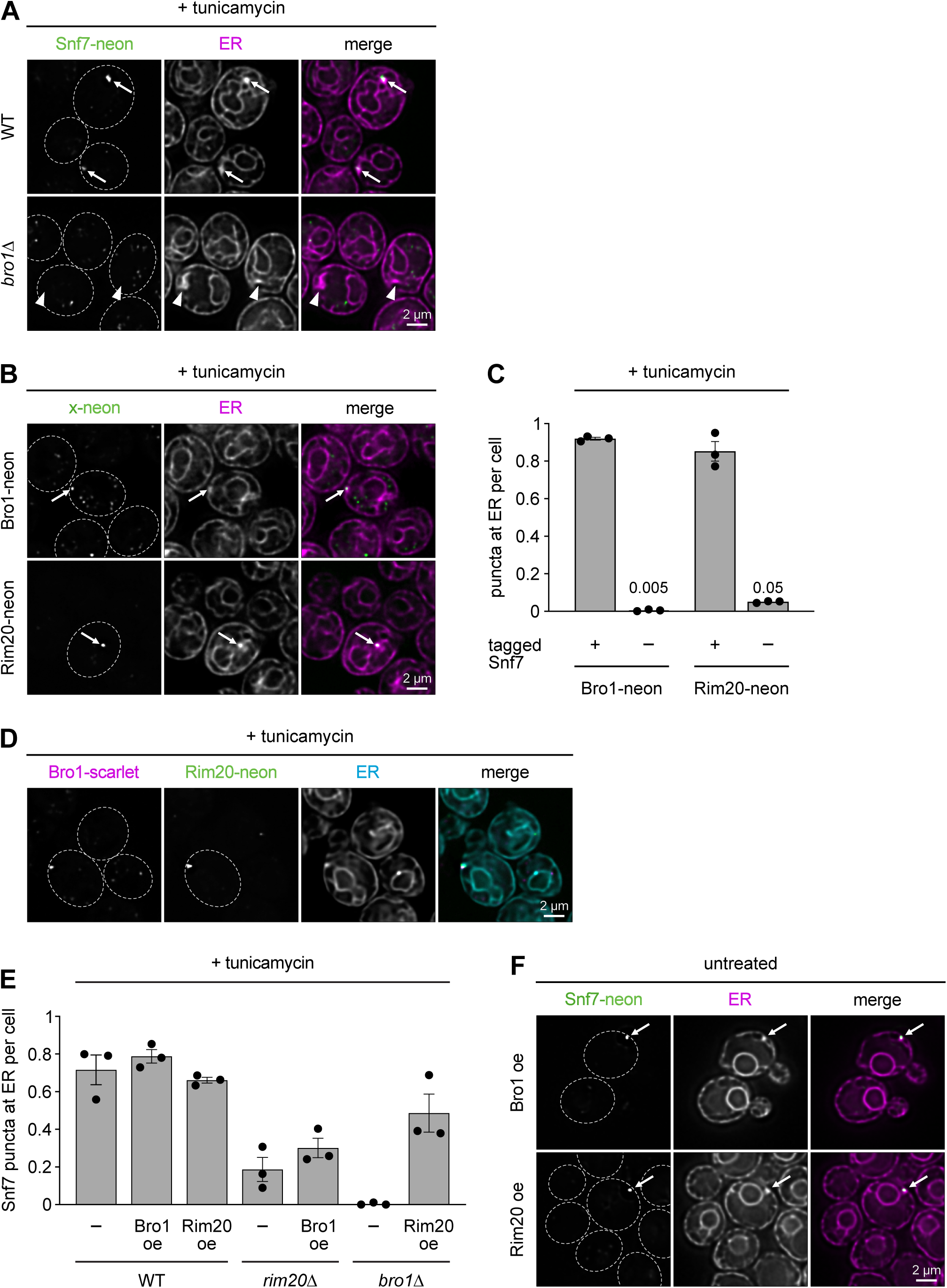
ER recruitment of Snf7 requires ESCRT-associated proteins Bro1 and Rim20. **(A)** Deconvolved fluorescent images of tunicamycin-treated wild-type (WT) and *bro1Δ* cells expressing Snf7-neon and the ER marker cherry-Ubc6. Dotted lines indicate cell boundaries. Arrows mark ER clusters co-localising with Snf7-neon puncta, arrowheads mark ER clusters not co-localising with Snf7-neon. **(B)** Deconvolved fluorescent images of tunicamycin-treated cells expressing Bro1-neon or Rim20-neon along with cherry-Ubc6. Dotted lines indicate cell boundaries. Arrows mark ER clusters co-localising with Bro1-neon or Rim20-neon puncta. The images show that ER-localised Bro1 and Rim20 puncta are present also in cells containing only endogenous Snf7. **(C)** Quantification of ER-localised Bro1-neon or Rim20-neon puncta per cell in tunicamycin-treated cells with or without additional expression of Snf7-scarlet (tagged Snf7). Data are the mean of n = 3 biological replicates, error bars indicate the standard error of the mean. **(D)** Deconvolved fluorescent images of tunicamycin-treated cells expressing Bro1-scarlet, Rim20-neon and the ER marker BFP-Ubc6. Dotted lines indicate cell boundaries. **(E)** Quantification of Snf7-neon puncta at ER clusters per cell in tunicamycin-treated and untreated wild-type (WT), *rim20Δ* and *bro1Δ* cells overexpressing Bro1 or Rim20 where indicated (oe = overexpression). Data are the mean of n = 3 biological replicates, error bars indicate the standard error of the mean. **(F)** Deconvolved fluorescent images of untreated wild-type cells expressing Snf7-neon and cherry-Ubc6, and overexpressing Bro1 or Rim20. Dotted lines indicate cell boundaries.

**Figure S3.**
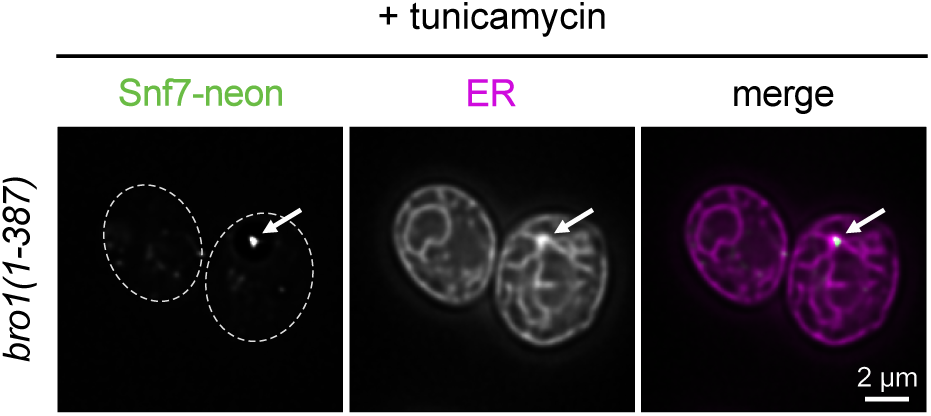
Snf7 recruitment to the ER requires specific features of Bro1. Deconvolved fluorescent images of tunicamycin-treated cells expressing Snf7-neon, the ER marker cherry-Ubc6 and truncated bro1(1-387) containing the Bro1 domain but lacking the V domain and proline-rich region. Dotted lines indicate cell boundaries. The Bro1 domain alone is sufficient to support Snf7 recruitment to the ER.

**Figure S4.**
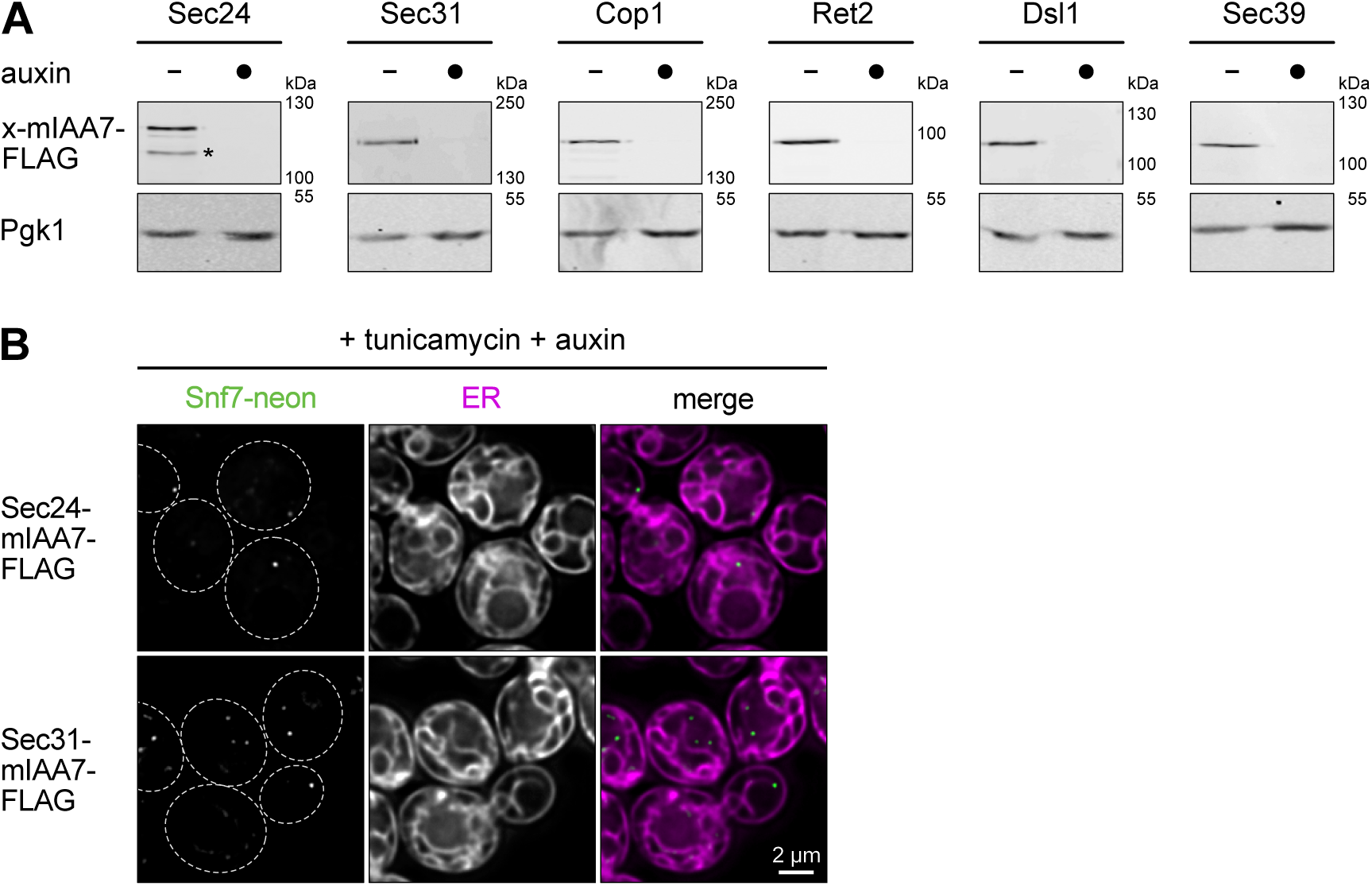
Auxin-induced degradation of COPII proteins, COPI proteins and Dsl1 complex subunits. **(A)** Western blot of FLAG tag from cells expressing Sec24, Sec31, Cop1, Ret2, Dsl1 or Sec39 tagged with the degron mIAA7-FLAG. Cells were treated with tunicamycin and, where indicated, additionally treated with auxin to induce target protein degradation. Pgk1 served as a loading control. The asterisk denotes a fragment of Sec24-mIAA7-FLAG, which presumably arose through degradation after cell lysis. **(B)** Deconvolved fluorescent images of cells expressing Snf7-neon, the ER marker cherry-Ubc6 and either Sec24-mIAA7-FLAG or Sec31-mIAA7-FLAG. Cells were treated with tunicamycin and additionally treated with auxin to induce Sec24 or Sec31 degradation. The ER in Sec24- and Sec31-depleted cells is disorganised and Snf7 only forms puncta at endosomes, not at the ER.

**Figure S5.**
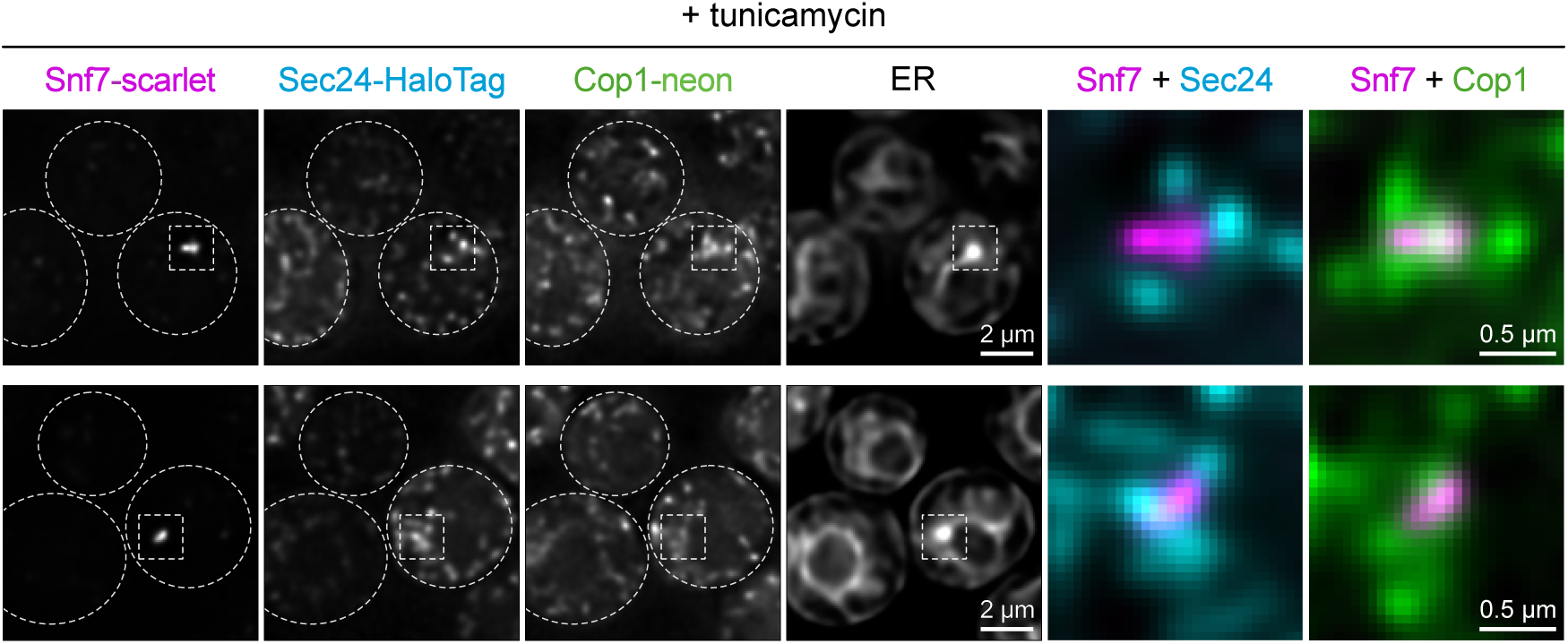
Snf7-containing ER subdomains are part of stress-induced ER-Golgi contacts. Deconvolved fluorescent images of tunicamycin-treated cells expressing Snf7-scarlet, the COPII protein Sec24-HaloTag, the COPI protein Cop1-neon and the ER marker BFP-Ubc6. The last two images show magnifications of the area marked with grey boxes in the first four images.

**Figure S6.**
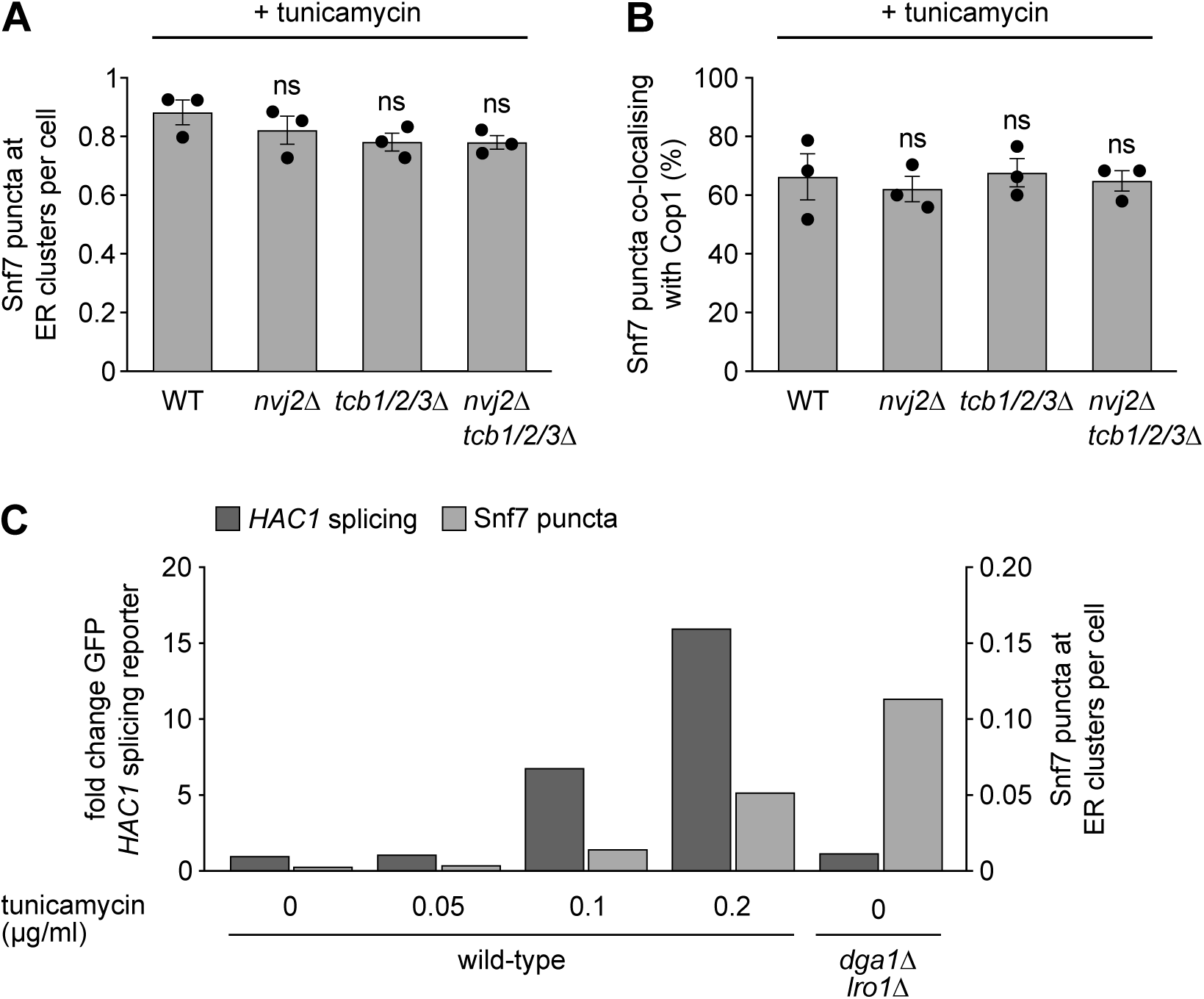
Stress-induced ER-Golgi contacts contain lipid transfer proteins. **(A)** Quantification of Snf7-neon puncta at ER clusters in tunicamycin-treated wild-type (WT) cells and the indicated mutants. Data are the mean of n = 3 biological replicates, error bars indicate the standard error of the mean. For each mutant, statistical significance of the difference to wild-type cells was evaluated with a two-tailed homoscedastic t-test. ns = not significant. **(B)** Quantification of co-localisation of Snf7-scarlet and the COPI protein Cop1-neon in tunicamycin-treated wild-type cells and the indicated mutants. Data are the mean of n = 3 biological replicates, error bars indicate the standard error of the mean. For each mutant, statistical significance of the difference to wild-type cells was evaluated with a two-tailed homoscedastic t-test. Ns = not significant. **(C)** Fold change GFP fluorescence arising from the *HAC1* splicing reporter (left y-axis) and quantification of Snf7-neon puncta at ER clusters per cell (right y-axis) in wild-type cells treated with different concentrations of tunicamycin and untreated *dga1*Δ *lro1*Δ cells. *HAC1* splicing was determined in cells expressing the *HAC1* reporter and Snf7 puncta were quantified in cells expressing Snf7-neon and the ER marker cherry-Ubc6. Data are from n = 1 experiment. *dga1Δ lro1Δ* cells show minimal activation of the unfolded protein response but more efficient recruitment of Snf7 to ER clusters than wild-type cells treated with up to 0.2 µg/ml tunicamycin.

**Table S4.** Plasmids used in this study.

| Plasmid | Alias | Source |
| --- | --- | --- |
| pFA6a-natNT2 | pSS032 | Janke, 2004 |
| pFA6a-hphNT1 | pSS033 | Janke, 2004 |
| pFA6a-kanMX6 | pSS034 | Longtine, 1998 |
| pFA6a-His3MX6 | pSS036 | Longtine, 1998 |
| pFA6a-kITRP1 | pSS876 | this study |
| pFA6a-LEU2 | pSS689 | this study |
| pFA6a-URA3 | pSS690 | this study |
| pFA6a-mCherry-HisMX6 | pSS063 | this study |
| pFA6a-mCherry-HDEL-hph | pCY3092-02 | Young, 2013 |
| pFA6a-mScarlet-i-kanMX6 | pSS917 | Papagiannidis, 2021 |
| pFA6a-mScarlet-i-natMX6 | pSS920 | this study |
| pFA6a-mNeonGreen-HA-HisMX6 | pSS694 | this study |
| pFA6a-mNeonGreen-HA-natNT2 | pSS695 | this study |
| pFA6a-mNeonGreen-HisMX6 | pSS447 | Papagiannidis, 2021 |
| pFA6a-mNeonGreen-natNT2 | pBB1 | Michael Knop |
| pFA6a-HaloTag-kanMX6 | pSS1409 | this study |
| pFA6a-hph-P<sub>TEF</sub> | pMaM215 | Michael Knop |
| pFA6a-nat-P<sub>ADH</sub>-yeGFP | pYM-N7 | Janke, 2004 |
| pFA6a-TurboID-3xmyc-kanMX6 | pSS1061 | Larochelle, 2019 |
| pFA6a-mIAA7-3FLAG-kanMX6 | pSS1154 | this study |
| pRS416-P<sub>VPS24</sub>-Snf7-LAP-mNeonGreen | pSS951 | Schäfer, 2020 |
| pRS416-P<sub>VPS24</sub>-Snf7-LAP-mScarlet-i | pSS1060 | this study |
| pRS415-P<sub>CYC</sub> | pSS021 | Mumberg, 1995 |
| pRS415-P<sub>VPS24</sub>-Snf7-LAP-mScarlet-i | pSS1570 | this study |
| pRS413-P<sub>CYC</sub> | pSS014 | Mumberg, 1995 |
| pRS413-P<sub>VPS24</sub>-Snf7-LAP-mScarlet-i | pSS1003 | this study |
| pRS416-Mup1-pHluorin | pLZ78 | Ivashov, 2020 |
| pRS404-P<sub>GPD</sub>-mCherry-Ubc6 | pSS962 | Schäfer, 2020 |
| pRS406-P<sub>TEF</sub>-TagBFP-Ubc6 | pSS988 | Platzek, 2025 |
| pRS304 | pSS003 | Sikorski, 1989 |
| pRS304-P<sub>TEF</sub>-TagBFP-Ubc6 | pSS1004 | this study |
| pRS404-P<sub>GPD</sub>-TagBFP-Ubc6 | pSS1310 | this study |
| pRS316-Ire1-HA | pPW1547 | Peter Walter |
| pRS315 | pSS008 | Sikorski, 1989 |
| pRS315-Ire1-HA | pSS320 | this study |
| pRS315-Ire1(mScarlet-i)-HA | pSS963 | this study |
| pRS316-GFP-Atg8 | pSS127 | Kirisako, 1999 |
| pRS413-P<sub>ADH</sub> | pSS015 | Mumberg, 1995 |
| pRS413-Bro1 | pSS964 | this study |
| pRS413-bro1(K246A) | pSS992 | this study |
| pRS413-bro1(Y320D) | pSS991 | this study |
| pRS303H-P<sub>GPD</sub>-TagBFP | pSS349 | Szoradi, 2018 |
| pRS303H-P<sub>GPD</sub>-TagBFP-PLCdelta-PH2 | pDK45 | Michael Knop |
| pNH605-Padh-OsTIR(F74G) | pSS1354 | Hubbe, 2026 |
| pRS305-HAC1-splicing-reporter | pDEP005 | Pincus, 2010 |

**Table S5.** Strains used in this study. BFP = TagBFP, Cherry = mCherry, halo = HaloTag, hph = hygromycin B resistance gene, kan = kanamycin resistance gene, nat = nourseothricin resistance gene, neon = mNeonGreen, scarlet = mScarlet-i.

| Strain | Relevant genotype |
| --- | --- |
| SSY122 | *ADE2 leu2-3,112 trp1-1 ura3-1 his3-11,15 MATa* |
| SSY2432 | *ura3::P<sub>VPS24</sub>-Snf7-neon-URA3* |
| SSY2583 | *ura3::P<sub>VPS24</sub>-Snf7-scarlet-URA3* |
| SSY795 | *his3::P<sub>GPD</sub>-BFP-hph* |
| SSY3677 | *his3::P<sub>GPD</sub>-BFP-hph ura3::Mup1-pHluorin-URA3* |
| SSY3669 | *his3::P<sub>GPD</sub>-BFP-hph ura3::Mup1-pHluorin-URA3 snf7Δ::nat* |
| SSY4375 | *his3::P<sub>GPD</sub>-BFP-hph ura3::Mup1-pHluorin-URA3 leu2::P<sub>VPS24</sub>-Snf7-scarlet-LEU2* |
| SSY4469 | *his3::P<sub>GPD</sub>-BFP-hph ura3::Mup1-pHluorin-URA3 leu2::P<sub>VPS24</sub>-Snf7-scarlet-LEU2 snf7Δ::nat* |
| SSY2544 | *ura3::P<sub>VPS24</sub>-Snf7-neon-URA3 trp1::P<sub>GPD</sub>-cherry-Ubc6-TRP1* |
| SSY3008 | *ura3::P<sub>VPS24</sub>-Snf7-neon-URA3 trp1::P<sub>GPD</sub>-cherry-Ubc6-TRP1 snf7Δ::nat* |
| SSY4319 | *ura3::P<sub>VPS24</sub>-Snf7-neon-URA3 KAR2-cherry-HDEL-hph* |
| SSY2710 | *ura3::P<sub>VPS24</sub>-Snf7-neon-URA3 RTN1-cherry::HIS3* |
| SSY4320 | *ura3::P<sub>VPS24</sub>-Snf7-neon-URA3 SEC63-scarlet::kan* |
| SSY4318 | *ura3::P<sub>VPS24</sub>-Snf7-neon-URA3 leu2::lre1-scarlet-LEU2* |
| SSY3607 | *trp1::P<sub>TEF</sub>-BFP-Ubc6-TRP1 his3::P<sub>VPS24</sub>-Snf7-scarlet-HIS3 pRS316-GFP-Atg8* |
| SSY2553 | *ura3::P<sub>VPS24</sub>-Snf7-neon-URA3 trp1::P<sub>GPD</sub>-cherry-Ubc6-TRP1 atg7Δ::nat* |
| SSY3785 | *ura3::P<sub>VPS24</sub>-Snf7-scarlet-URA3 trp1::P<sub>GPD</sub>-BFP-Ubc6-TRP1 SEC24-neon-HA-HIS3 chm7Δ::nat* |
| SSY2555 | *ura3::P<sub>VPS24</sub>-Snf7-neon-URA3 trp1::P<sub>GPD</sub>-cherry-Ubc6-TRP1 bro1Δ::nat* |
| SSY4314 | *ura3::P<sub>VPS24</sub>-Snf7-neon-URA3 trp1::P<sub>GPD</sub>-cherry-Ubc6-TRP1 vps27Δ::nat hse1Δ::hph* |
| SSY2549 | *ura3::P<sub>VPS24</sub>-Snf7-neon-URA3 trp1::P<sub>GPD</sub>-cherry-Ubc6-TRP1 vps23Δ::kan* |
| SSY2557 | *ura3::P<sub>VPS24</sub>-Snf7-neon-URA3 trp1::P<sub>GPD</sub>-cherry-Ubc6-TRP1 vps36Δ::nat* |
| SSY2546 | *ura3::P<sub>VPS24</sub>-Snf7-neon-URA3 trp1::P<sub>GPD</sub>-cherry-Ubc6-TRP1 vps20Δ::nat* |
| SSY2556 | *ura3::P<sub>VPS24</sub>-Snf7-neon-URA3 trp1::P<sub>GPD</sub>-cherry-Ubc6-TRP1 chm7Δ::nat* |
| SSY3040 | *ura3::P<sub>VPS24</sub>-Snf7-neon-URA3 trp1::P<sub>GPD</sub>-cherry-Ubc6-TRP1 rim20Δ::nat* |
| SSY3039 | *ura3::P<sub>VPS24</sub>-Snf7-neon-URA3 trp1::P<sub>GPD</sub>-cherry-Ubc6-TRP1 ygr122wΔ::nat* |
| SSY2834 | *ura3::P<sub>VPS24</sub>-Snf7-scarlet-URA3 trp1::P<sub>GPD</sub>-BFP-Ubc6-TRP1 BRO1-neon-nat* |
| SSY3073 | *ura3::P<sub>VPS24</sub>-Snf7-scarlet-URA3 trp1::P<sub>GPD</sub>-BFP-Ubc6-TRP1 RIM20-neon-HA-HIS3* |
| SSY2693 | *ura3::P<sub>TEF</sub>-cherry-Ubc6-URA3 BRO1-neon-HA-nat* |
| SSY3130 | *trp1::P<sub>GPD</sub>-cherry-Ubc6-TRP1 RIM20-neon-HA-HIS3* |
| SSY3267 | *ura3::P<sub>TEF</sub>-BFP-Ubc6-URA3 RIM20-neon-HA-HIS3 BRO1-scarlet-nat* |
| SSY3556 | *ura3::P<sub>VPS24</sub>-Snf7-neon-URA3 trp1::P<sub>GPD</sub>-cherry-Ubc6-TRP1 hph-P<sub>TEF</sub>-BRO1* |
| SSY3557 | *ura3::P<sub>VPS24</sub>-Snf7-neon-URA3 trp1::P<sub>GPD</sub>-cherry-Ubc6-TRP1 hph-P<sub>TEF</sub>-RIM20* |
| SSY3558 | *ura3::P<sub>VPS24</sub>-Snf7-neon-URA3 trp1::P<sub>GPD</sub>-cherry-Ubc6-TRP1 hph-P<sub>TEF</sub>-BRO1 rim20Δ::nat* |
| SSY3559 | *ura3::P<sub>VPS24</sub>-Snf7-neon-URA3 trp1::P<sub>GPD</sub>-cherry-Ubc6-TRP1 hph-P<sub>TEF</sub>-RIM20 bro1Δ::nat* |
| SSY4470 | *ura3::P<sub>VPS24</sub>-Snf7-neon-URA3 trp1::P<sub>GPD</sub>-cherry-Ubc6-TRP1 bro1(1-387)::nat* |
| SSY3707 | *his3::P<sub>GPD</sub>-BFP-hph ura3::Mup1-pHluorin-URA3 bro1Δ::nat* |
| SSY3055 | *bro1(K246A)* |
| SSY3466 | *bro1(Y320D)* |
| SSY3735 | *bro1(K246A) his3::P<sub>GPD</sub>-BFP-hph ura3::Mup1-pHluorin-URA3* |
| SSY3733 | *bro1(Y320D) his3::P<sub>GPD</sub>-BFP-hph ura3::Mup1-pHluorin-URA3* |
| SSY4324 | *bro1(K246A) ura3::P<sub>VPS24</sub>-Snf7-neon-URA3 trp1::P<sub>GPD</sub>-cherry-Ubc6-TRP1* |
| SSY4323 | *bro1(Y320D) ura3::P<sub>VPS24</sub>-Snf7-neon-URA3 trp1::P<sub>GPD</sub>-cherry-Ubc6-TRP1* |
| SSY3474 | *bro1(Y320D)-neon-HA::HIS3 trp1::P<sub>TEF</sub>-BFP-Ubc6-TRP1 ura3::P<sub>VPS24</sub>-Snf7-scarlet-URA3* |
| SSY3378 | *bro1(K246A)-neon-HA::HIS3 trp1::P<sub>TEF</sub>-BFP-Ubc6-TRP1 ura3::P<sub>VPS24</sub>-Snf7-scarlet-URA3* |
| SSY2969 | *ura3::P<sub>VPS24</sub>-Snf7-neon-URA3 BRO1-TurboID-3xmyc-kan vps27Δ::nat* |
| SSY3070 | *bro1(K246A)-TurboID-3xmyc-kan ura3::P<sub>VPS24</sub>-Snf7-neon-URA3 vps27Δ::nat* |
| SSY4334 | *bro1(Y320D)-TurboID-3xmyc-kan ura3::P<sub>VPS24</sub>-Snf7-neon-URA3 vps27Δ::nat* |
| SSY3290 | *ura3::P<sub>VPS24</sub>-Snf7-neon-URA3 BRO1-TurboID-3xmyc-kan vps27Δ::nat pep4Δ::hph prb1Δ::TRP1 chm7Δ::HIS3* |
| SSY3292 | *ura3::P<sub>VPS24</sub>-Snf7-neon-URA3 bro1(K246A)-TurboID-3xmyc-kan vps27Δ::nat pep4Δ::hph prb1Δ::TRP1 chm7Δ::HIS3* |
| SSY3005 | *ura3::P<sub>VPS24</sub>-Snf7-scarlet-URA3 trp1::P<sub>GPD</sub>-BFP-Ubc6-TRP1 IST1-neon-HA-HIS3* |
| Y8205 | *can1Δ::P<sub>STE2</sub>-Sp_his5 lyp1Δ::P<sub>STE3</sub>-LEU2 his3Δ1 leu2Δ0 ura3Δ0 MATα* |
| SSY2643 | *Y8205 his3::P<sub>GPD</sub>-BFP-PLCdelta-PH2-hph ura3::P<sub>VPS24</sub>-Snf7-neon-ura3::P<sub>TEF</sub>-cherry-Ubc6-nat vps27Δ::URA3* |
| SSY2644 | *Y8205 his3::P<sub>GPD</sub>-BFP-PLCdelta-PH2-hph ura3::P<sub>VPS24</sub>-Snf7-neon-ura3::P<sub>TEF</sub>-cherry-Ubc6-nat vps27Δ::URA3 bro1Δ::kan* |
| SSY3361 | *ura3::P<sub>VPS24</sub>-Snf7-neon-URA3 trp1::P<sub>GPD</sub>-cherry-Ubc6-TRP1 ted1Δ::HIS3* |
| SSY3359 | *ura3::P<sub>VPS24</sub>-Snf7-neon-URA3 trp1::P<sub>GPD</sub>-cherry-Ubc6-TRP1 bst1Δ::HIS3* |
| SSY2832 | *ura3::P<sub>VPS24</sub>-Snf7-neon-URA3 trp1::P<sub>GPD</sub>-cherry-Ubc6-TRP1 erv25Δ::nat* |
| SSY2833 | *ura3::P<sub>VPS24</sub>-Snf7-neon-URA3 trp1::P<sub>GPD</sub>-cherry-Ubc6-TRP1 emp24Δ::nat* |
| SSY4856 | *ura3::P<sub>VPS24</sub>-Snf7-neon-URA3 trp1::P<sub>GPD</sub>-cherry-Ubc6-TRP1 tsc3Δ::nat* |
| SSY2936 | *ura3::P<sub>VPS24</sub>-Snf7-scarlet-URA3 trp1::P<sub>GPD</sub>-BFP-Ubc6-TRP1 EMP24-neon-HIS3* |
| SSY3358 | *ura3::P<sub>VPS24</sub>-Snf7-scarlet-URA3 trp1::P<sub>GPD</sub>-BFP-Ubc6-TRP1 emp24(F196A,F197A)-neon-HA-HIS3* |
| SSY4717 | *ura3::P<sub>VPS24</sub>-Snf7-neon-URA3 trp1::P<sub>GPD</sub>-cherry-Ubc6-TRP1 SEC24-mIAA7-FLAG-kan leu2::P<sub>ADH</sub>-OsTIR1(F74G)-LEU2* |
| SSY4718 | *ura3::P<sub>VPS24</sub>-Snf7-neon-URA3 trp1::P<sub>GPD</sub>-cherry-Ubc6-TRP1 SEC31-mIAA7-FLAG-kan leu2::P<sub>ADH</sub>-OsTIR1(F74G)-LEU2* |
| SSY3990 | *ura3::P<sub>VPS24</sub>-Snf7-neon-URA3 trp1::P<sub>GPD</sub>-cherry-Ubc6-TRP1 COP1-mIAA7-FLAG-kan leu2::P<sub>ADH</sub>-OsTIR1(F74G)-LEU2* |
| SSY3991 | *ura3::P<sub>VPS24</sub>-Snf7-neon-URA3 trp1::P<sub>GPD</sub>-cherry-Ubc6-TRP1 RET2-mIAA7-FLAG-kan leu2::P<sub>ADH</sub>-OsTIR1(F74G)-LEU2* |
| SSY4721 | *ura3::P<sub>VPS24</sub>-Snf7-neon-URA3 trp1::P<sub>GPD</sub>-cherry-Ubc6-TRP1 DSL1-mIAA7-FLAG-kan leu2::P<sub>ADH</sub>-OsTIR1(F74G)-LEU2* |
| SSY4719 | *ura3::P<sub>VPS24</sub>-Snf7-neon-URA3 trp1::P<sub>GPD</sub>-cherry-Ubc6-TRP1 SEC39-mIAA7-FLAG-kan leu2::P<sub>ADH</sub>-OsTIR1(F74G)-LEU2* |
| SSY3897 | *ura3::P<sub>VPS24</sub>-Snf7-scarlet-URA3 trp1::P<sub>GPD</sub>-BFP-Ubc6-TRP1 COP1-neon-HA-HIS3 SEC24-halo-kan* |
| SSY3493 | *ura3::P<sub>VPS24</sub>-Snf7-scarlet-URA3 trp1::P<sub>GPD</sub>-BFP-Ubc6-TRP1 COP1-neon-HA-HIS3* |
| SSY3555 | *ura3::P<sub>VPS24</sub>-Snf7-scarlet-URA3 trp1::P<sub>GPD</sub>-BFP-Ubc6-TRP1 RET2-neon-HA-HIS3* |
| SSY4609 | *ura3::P<sub>VPS24</sub>-Snf7-scarlet-URA3 trp1::P<sub>GPD</sub>-BFP-Ubc6-TRP1 SEC21-neon-HIS3* |
| SSY4610 | *ura3::P<sub>VPS24</sub>-Snf7-scarlet-URA3 trp1::P<sub>GPD</sub>-BFP-Ubc6-TRP1 SEC27-neon-HIS3* |
| SSY4611 | *ura3::P<sub>VPS24</sub>-Snf7-scarlet-URA3 trp1::P<sub>GPD</sub>-BFP-Ubc6-TRP1 SEC28-neon-HIS3* |
| SSY4170 | *ura3::P<sub>VPS24</sub>-Snf7-scarlet-URA3 trp1::P<sub>GPD</sub>-BFP-Ubc6-TRP1 GRH1-neon-HIS3* |
| SSY4213 | *ura3::P<sub>VPS24</sub>-Snf7-scarlet-URA3 trp1::P<sub>GPD</sub>-BFP-Ubc6-TRP1 MNN9-neon-HIS3* |
| SSY4467 | *ura3::P<sub>VPS24</sub>-Snf7-scarlet-URA3 trp1::P<sub>GPD</sub>-BFP-Ubc6-TRP1 COP1-halo-kan AUR1-neon-HIS* |
| SSY4174 | *ura3::P<sub>VPS24</sub>-Snf7-scarlet-URA3 trp1::P<sub>GPD</sub>-BFP-Ubc6-TRP1 nat-P<sub>ADH</sub>-yeGFP-GOS1* |
| SSY4175 | *ura3::P<sub>VPS24</sub>-Snf7-scarlet-URA3 trp1::P<sub>GPD</sub>-BFP-Ubc6-TRP1 GGA2-neon-HIS3* |
| SSY4716 | *ura3::P<sub>VPS24</sub>-Snf7-scarlet-URA3 trp1::P<sub>GPD</sub>-BFP-Ubc6-TRP1 APL6-neon-HIS3* |
| SSY4373 | *ura3::P<sub>VPS24</sub>-Snf7-scarlet-URA3 trp1::P<sub>GPD</sub>-BFP-Ubc6-TRP1 DSL1-neon-HA-HIS3 COP1-halo-kan* |
| SSY4214 | *ura3::P<sub>VPS24</sub>-Snf7-scarlet-URA3 trp1::P<sub>GPD</sub>-BFP-Ubc6-TRP1 GYP1-neon-HIS3* |
| SSY4938 | *ura3::P<sub>VPS24</sub>-Snf7-scarlet-URA3 trp1::P<sub>GPD</sub>-BFP-Ubc6-TRP1 ARF2-neon-HIS3* |
| SSY3986 | *ura3::P<sub>VPS24</sub>-Snf7-scarlet-URA3 trp1::P<sub>GPD</sub>-BFP-Ubc6-TRP1 NVJ2-neon-HIS3* |
| SSY4153 | *ura3::P<sub>VPS24</sub>-Snf7-scarlet-URA3 trp1::P<sub>GPD</sub>-BFP-Ubc6-TRP1 TCB3-neon-HIS3* |
| SSY4376 | *ura3::P<sub>VPS24</sub>-Snf7-scarlet-URA3 trp1::P<sub>GPD</sub>-BFP-Ubc6-TRP1 TCB3-neon-HIS3 COP1-halo-kan* |
| SSY4190 | *bro1(Y320D) trp1::P<sub>TEF</sub>-BFP-Ubc6-TRP1 ura3::P<sub>VPS24</sub>-Snf7-scarlet-URA3 TCB3-neon-HIS3* |
| SSY3987 | *ura3::P<sub>VPS24</sub>-Snf7-neon-URA3 trp1::P<sub>GPD</sub>-cherry-Ubc6-TRP1 nvj2Δ::nat* |
| SSY4200 | *ura3::P<sub>VPS24</sub>-Snf7-neon-URA3 trp1::P<sub>GPD</sub>-cherry-Ubc6-TRP1 tcb1Δ::LEU2 tcb2Δ::nat tcb3Δ::hph* |
| SSY4725 | *ura3::P<sub>VPS24</sub>-Snf7-neon-URA3 trp1::P<sub>GPD</sub>-cherry-Ubc6-TRP1 tcb1Δ::LEU2 tcb2Δ::nat tcb3Δ::hph nvj2Δ::kan* |
| SSY4747 | *ura3::P<sub>VPS24</sub>-Snf7-scarlet-URA3 trp1::P<sub>GPD</sub>-BFP-Ubc6-TRP1 nvj2Δ::kan COP1-neon-HA-HIS3* |
| SSY4708 | *ura3::P<sub>VPS24</sub>-Snf7-scarlet-URA3 trp1::P<sub>GPD</sub>-BFP-Ubc6-TRP1 tcb1Δ::LEU2 tcb2Δ::nat tcb3Δ::hph COP1-neon-HA-HIS3* |
| SSY4709 | *ura3::P<sub>VPS24</sub>-Snf7-scarlet-URA3 trp1::P<sub>GPD</sub>-BFP-Ubc6-TRP1 tcb1Δ::LEU2 tcb2Δ::nat tcb3Δ::hph nvj2Δ::kan COP1-neon-HA-HIS3* |
| SSY4855 | *lro1Δ::hph dga1Δ::kan* |
| SSY4309 | *bro1(Y320D) dga1Δ::kan lro1Δ::hph* |
| SSY4189 | *ura3::P<sub>VPS24</sub>-Snf7-neon-URA3 trp1::P<sub>GPD</sub>-cherry-Ubc6-TRP1 dga1Δ::HIS3 lro1Δ::nat* |
| SSY4545 | *leu2::HAC1-splicing reporter-LEU2* |
| SSY5146 | *leu2::HAC1-splicing reporter-LEU2 lro1Δ::kan* |
| SSY5147 | *leu2::HAC1-splicing reporter-LEU2 dga1Δ::nat* |
| SSY5148 | *leu2::HAC1-splicing reporter-LEU2 lro1Δ::kan dga1Δ::nat* |

**Table S6.** Milling parameters used with the AutoLamella package. All units are in meter, except for the milling current, which is in ampere.

| parameters | micro expansions | parameters | milling step 1 | milling step 2 | milling step 3 |
| --- | --- | --- | --- | --- | --- |
| width | 1.44e-6 | lamella width | 12.0e-6 | 11.0e-6 | 10.5e-6 |
| height | 12.83e-6 | lamella height | 0.5e-6 | 0.5e-6 | 0.5e-6 |
| distance | 4.33e-6 | trench height | 5.0e-6 | 1.4e-6 | 1.0e-6 |
| milling current | 1.0e-9 | depth | 0.9e-6 | 0.9e-6 | 0.5 e-6 |
|  |  | offset | 1.0e-6 | 0.5e-6 | 0.15 e-6 |
|  |  | milling current | 1.0e-9 | 0.3e-9 | 0.1e-9 |

